# The Multisensory Correlation Detector model explains temporal integration in the Rubber Hand Illusion

**DOI:** 10.64898/2026.11.16.766969

**Authors:** Piotr Litwin

## Abstract

The Rubber Hand Illusion (RHI) demonstrates that body ownership is flexibly constructed in real-time. Since RHI occurrence patterns strongly adhere to key principles of multisensory integration, the illusion can be modeled computationally as an optimal cue integration process. Here, we model RHI occurrence probabilities under different asynchrony conditions as a function of temporal visuo-tactile correlations, using the Multisensory Correlation Detector (MCD) model. Three computational simulations using human data from two psychophysical RHI detection experiments were carried out. The results of Simulation 1 show that the MCD accounts for observed probabilities in RHI detection tasks, fitting the empirical data and predicting out-of-sample probabilities more closely than a Bayesian Causal Inference model. Simulation 2 demonstrates that MCD predicts visuo-tactile stimulation patterns with complex temporal structures should lead to higher RHI occurrence probabilities, offering the first computational mechanism for previously demonstrated effect of stimulation complexity. Finally, Simulation 3 shows that MCD relates skewed distributions of RHI occurrence to asymmetrical time constants in early filtering of visual and tactile signals, suggesting a common mechanism for simultaneity and body ownership perception. These findings indicate that MCD is a promising alternative for advancing our understanding of computational mechanisms underlying body ownership, with several unique strengths.

## Introduction

The sense of body ownership—our implicit understanding of what constitutes our physical self—emerges from integration of multisensory inputs (Botvinick, 2004; Ehrsson, 2012; Kilteni et al., 2015; Makin et al., 2008). A coherent representation of our body is continuously constructed by combining multiple sensory signals in real time; mainly visual, tactile, and proprioceptive signals (Litwin, 2020), as well as potentially other, less directly related inputs from the auditory (Radziun & Ehrsson, 2018), interoceptive (Suzuki et al., 2013), and vestibular (Preuss & Ehrsson, 2019) domains. The malleability of this process is strikingly demonstrated by the Rubber Hand Illusion (RHI), where the synchronous stroking of a visible fake hand and one’s hidden real hand can induce a sense of ownership over the fake limb. Since it was introduced by Botvinick and Cohen (1998), the RHI has become a cornerstone paradigm for investigating body ownership (Riemer et al., 2019). RHI occurrence and strength depend on factors such as synchrony (Bekrater-Bodmann et al., 2014; Shimada et al., 2009), the presence and complexity of visuo-tactile stimulation (Litwin et al., 2020), the distance between the two hands (Kalckert & Ehrsson, 2014; Kalckert et al., 2019), as well as sensory noise levels (Chancel & Ehrsson, 2023; Chancel et al., 2022a), with effects consistently in the direction predicted by multisensory integration theory. These correspondences suggest that mechanisms underlying body ownership likely rely on statistically optimal integration (Ehrsson, 2020).

Given the strong adherence to key multisensory integration principles—including spatial and temporal proximity as well as inverse effectiveness rules—the RHI lends itself to computational modeling. Current computational approaches include Bayesian (Chancel & Ehrsson, 2023; Chancel et al., 2022a, b; Fang et al., 2019; Samad et al., 2015, 2018), predictive coding (Hinz et al., 2018), active inference (Lanillos & Cheng, 2018; Lanillos et al., 2021; Rood et al., 2020), and deterministic chaotic network models (Ponzi & Suzuki, 2025). The Bayesian framework is currently at the forefront in terms of explanatory scope. Bayesian models frame body ownership as a computational problem of causal inference, in which the brain determines whether visual and somatosensory signals arise from a common source. The tactile receptor surface is co-extensive with the skin; therefore, from the brain’s perspective, tactile signals can only originate from the contact of one’s body with the world (Litwin, 2020). As a result, inference of a common cause inevitably leads to the transfer of ownership to the location where the touch is clearly seen: the rubber hand. The strength of Bayesian models lies in formalizing how bottom-up sensory evidence can be combined with top-down priors—with respect to the reliability of the former—to determine whether visual and tactile signals share a common origin. These models have successfully captured general RHI occurrence patterns, correctly predicting a decreased probability of the illusion with increased spatial (Fang et al., 2019; Samad et al., 2015, 2018) and temporal (Chancel & Ehrsson, 2023; Chancel et al., 2022a) disparity between visual and somatosensory signals, as well as an increased probability of RHI occurrence under conditions of elevated visual (Chancel et al., 2022a) and proprioceptive (Chancel & Ehrsson, 2023) noise.

A recent study by Chancel et al. (2022a) stands out as marking a significant advancement in Bayesian models for body ownership. The authors designed a psychophysical task in which participants observed automated taps delivered to the rubber hand in spatial congruence with their occluded real hand. The manipulation included trial-to-trial variations in tap synchrony and the level of visual noise. In each trial, participants reported whether the rubber hand felt like their own or not. A Bayesian Causal Inference (BCI) model, consisting of parameters related to the prior probability of a common cause, distinct sensory noise parameters for each noise condition, and the lapse rate, was fitted to observed RHI probabilities under different asynchrony and noise conditions. The BCI model provided an excellent fit to the data, accounting for the lower proportion of affirmative responses with increased asynchrony as well as the higher proportion of affirmative responses in conditions with higher visual noise. Chancel and Ehrsson (2023) used an analogous task with trial-to-trial variations in asynchrony and proprioceptive noise, showing that proprioceptive uncertainty also promotes the RHI, in line with BCI predictions. These findings were corroborated by an fMRI study, which showed that neural activity in the posterior parietal cortex closely fit BCI-predicted probabilities of RHI emergence (Chancel et al., 2022b). Similarly, in macaque monkeys, premotor cortex activity reflects the posterior probability of a common cause as spatial disparity between visual and proprioceptive signals varies (Fang et al., 2019). Empirical evidence converging across various neuroimaging methods and studies suggests a possible neural implementation of causal inference for body ownership (D’Angelo et al., 2026).

Despite these advancements, Bayesian models of the RHI face some important limitations. Most importantly, BCI models are not stimulus-computable: they do not operate on raw sensory signals, but instead handle abstract, experimenter-defined parameters. This model structure lacks the flexibility to accommodate certain stimulus-driven factors that might play important roles in constituting body ownership, such as the structural temporal complexity of visuo-tactile signals (Litwin, 2020). Relatedly, in BCI, individual differences can only arise from participant-specific values of these abstract parameters rather than from idiosyncrasies in raw signal processing. These limitations highlight the need for alternative approaches that can address these issues while maintaining the explanatory power of a Bayesian-optimal estimator, but without the computational strain that it entails.

The Multisensory Correlation Detector (MCD) model—initially proposed for insect motion perception (Hassenstein & Reichardt, 1956) and extrapolated to human motion perception (Adelson & Bergen, 1985)—has only recently been developed for multisensory integration (Parise & Ernst, 2016; Parise, 2025; Parise & Ernst, 2025). The MCD model processes two signals from different sensory modalities through a unified processing unit (Figure 1). Unisensory signals are initially filtered with either low-pass filters that pass low-frequency components, encoding stimulus intensity while being less sensitive to rapid changes (original implementation; Parise & Ernst, 2016) or band-pass filters tuned to specific frequency ranges in order to detect intensity changes (revised version; Parise & Ernst, 2025). Filtered signals represent the lumped impulse response of each signal after transduction, transmission and early unisensory processing (Parise & Ernst, 2016, p. 2). Signals are then fed into two cross-wired correlation detectors which multiply the signals after introducing a temporal shift to one of them through a late-stage low-pass filter, tuning themselves to different temporal orders of the signals. Following this late-stage filtering, the signals are either multiplied or subtracted, generating time-varying responses of the whole unit that represent the local temporal correlation and lag between the signals. These time-specific responses can be averaged over the analyzed time period to produce general indices of temporal correlation and lag.

**Figure 1.**
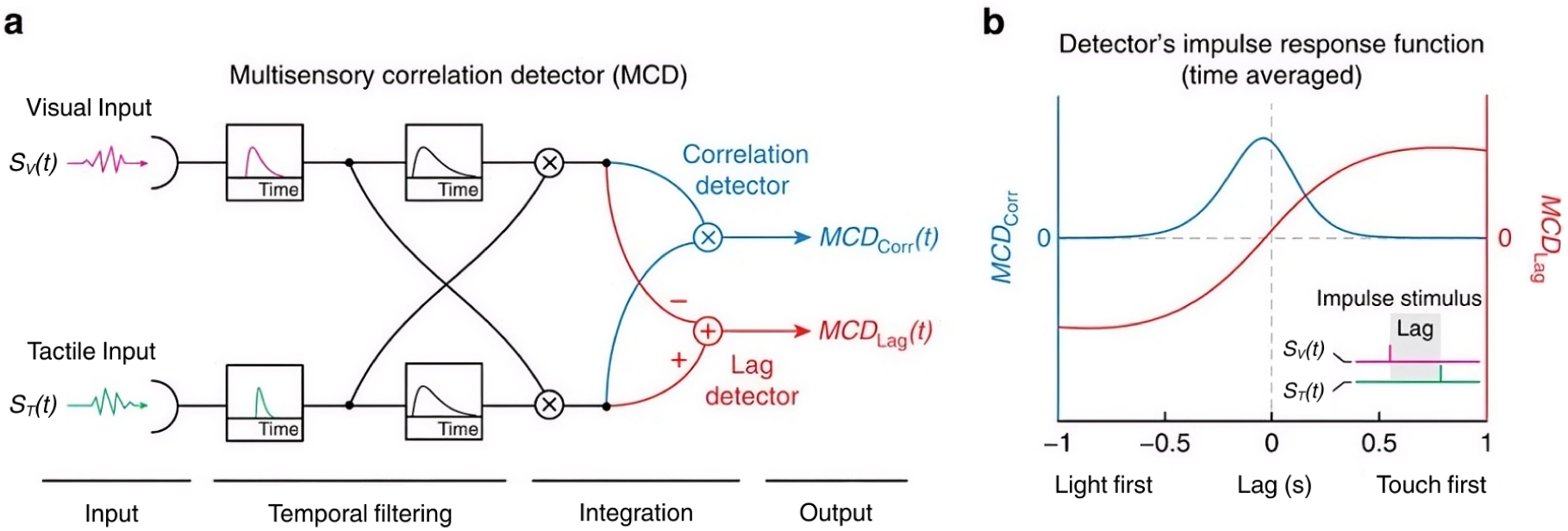
Multisensory Correlation Detector model for multisensory integration. (A) An example of MCD model implementation for visuo-tactile integration. The temporal filtering stage consists of two phases: early and late filtering. Small black dots indicate the stages at which signals enter into linear operations: multiplication (indicated by crosses) and subtraction (the "+" in the figure denotes the operation −S1 + S2, which is equivalent to S2 − S1). (B) The MCD model output (averaged over the signal duration) for an example visuo-tactile stream presented as a function of the offset between visual and tactile signals. The *MCD_corr_* indicates the correlation of signals and takes only positive values, while the *MCD_lag_* can take both positive and negative values, depending on which signal was detected as being presented first. The figure was adapted from Parise and Ernst (2016), published under a CC BY 4.0 license, and slightly modified for the context of visuo-tactile integration.

The MCD model offers a simple alternative to the causal inference problem: signals with highly correlated temporal structures are more likely to be inferred to stem from a common cause (cf. Parise et al., 2012) and thus be integrated. Although MCD lacks a dedicated Bayesian architecture, it demonstrated Bayes-optimal performance across various audiovisual tasks involving simultaneity and temporal order detection (Parise & Ernst, 2016, 2025). Another key advantage of MCD is that it operates directly on sensory input. Crucially, this feature is shared with biological cognitive systems which also operate directly on sensory signals. As a stimulus-computable model, MCD is capable of capturing effects that arise directly from relevant signal characteristics, such as structural complexity or the amount of noise. These signal properties do not have to be fed into the model as abstract parameters, as they are directly present in the signal itself, which renders MCD well suited for dynamic sensory data (Pesnot Lerousseau et al., 2022). Another important advantage of MCD is that it includes simple computations that are plausible to be carried out in the brain for real-time sensory processing (Parise & Ernst, 2016), which indicates its potential to link behavioral observations to candidate neural mechanisms. The model is biologically inspired and its filtering stages are considered a viable element of the human sensory processing system. For example, models assuming early low-pass filtering in the visual system provide accurate descriptions of the response characteristics of cones (Donner, 2021).

These key characteristics and strong mechanistic pedigree (Borst & Helmstaedter, 2015; Hassenstein & Reichardt, 1956) suggest the MCD’s potential to explain both behavioral observations and neural mechanisms of visuo-tactile integration for body ownership. In the current paper, we applied MCD to model RHI occurrence probabilities in a yes/no RHI detection task (Chancel & Ehrsson, 2023; Chancel et al., 2022a). Across three simulations, we demonstrate the capacity of the MCD model to explain various aspects of the illusion. First, we assessed how well the MCD model fits RHI detection probability data from two independent datasets under conditions without sensory noise, and how well it predicts RHI detection probabilities at asynchrony levels omitted from fitting. Ownership responses were either read out from the *MCD_corr_* output, under the assumption that the probability of ownership increases with the temporal correlation between visual and tactile signals (MCD-C), or jointly from *MCD_corr_*and the temporal order information carried by *MCD_lag_* (MCD-C+L). The results showed that the MCD fitted the data more closely than a BCI model restricted to the no-noise condition (BCI-0) and, in leave-one-asynchrony-out cross-validation, predicted the omitted levels better than BCI-0. The second simulation showed that the MCD model predicts higher rates of RHI occurrence for visuo-tactile stimulation patterns with complex (stochastic) temporal structures compared to regular (rhythmic) ones, which is in line with previous empirical findings (Litwin et al., 2020). This effect arises already at the level of the correlation detection mechanism rather than in the decisional read-out. Finally, the third simulation demonstrated that independently varying time constants of unisensory filters shifts the *point of strongest ownership* (PSO), while varying them jointly changes the width of the window *of subjective ownership* (WSO). This shows that MCD links individual differences in susceptibility to RHI under different asynchrony conditions to processing times of visuo-tactile signals. In turn, this connects RHI to other easily measurable psychophysical effects, such as visuo-tactile temporal binding window, which has been shown to be predictive of RHI susceptibility under moderate asynchrony conditions (Costantini et al., 2016). Together, these results indicate that MCD not only fits RHI detection data and predicts them out of sample, but also accounts for effects of signal complexity and visuo-tactile processing that are not captured by current BCI implementations.

## Materials and Methods

### Data

All models were fitted separately to two independent datasets collected with the same RHI detection paradigm. The first dataset (Chancel et al., 2022a) included a total of 3,780 yes/no responses from 15 participants who performed a detection task related to the subjective experience of the RHI (illusion present/absent). In each trial, participants received visuo-tactile stimulation in the form of spatially congruent taps on both hands. Participants were tapped on their index finger six times within a 12-second interval, after which they reported whether they had experienced the RHI. The experiment manipulated two factors: (1) the degree of visuo-tactile asynchrony (-500, -300, -150, 0, 150, 300, 500 ms offset, with negative values indicating that participants first observed the tap on the rubber hand and then felt it on their own hand) and (2) the degree of visual noise implemented using augmented reality glasses (0%, 30%, 50%). Each of the 21 conditions, resulting from all possible combinations of asynchrony and noise, was repeated 12 times, yielding a total of 252 trials per participant. For model fitting, we used only 1,260 trials from the no-noise condition, because the input to the MCD was defined in the temporal domain simply as presence or absence of a tap, and spatial noise could not be simulated in this one-dimensional signal. RHI occurrence probability was calculated across participants as the mean proportion of affirmative responses at each asynchrony condition. No participant-wise analyses were performed.

The second dataset was collected by Chancel and Ehrsson (2023). The only major difference between the studies concerned the noise condition, as the latter study manipulated proprioceptive rather than visual noise. This difference was not relevant, since the no-noise condition and the set of seven visuo-tactile asynchrony levels overlapped between the studies. The second dataset included 6,944 yes/no trials from 31 participants, of which 3,472 trials from the no-noise condition (16 repetitions of each asynchrony level per participant) were used for model fitting. Again, RHI occurrence probability was calculated across participants as the mean proportion of affirmative responses for seven different asynchrony levels, and no participant-wise analyses were made. In both studies, all participants provided written informed consent, and the experiments were approved by the Swedish Ethical Review Authority as stated in original publications (Chancel & Ehrsson, 2023; Chancel et al., 2022a).

#### Visuo-Tactile Signals

Visuo-tactile signal inputs for the MCD model were simulated as varying tapping patterns, differing in terms of temporal structure (regularity) and visuo-tactile asynchrony. The same seven asynchrony levels as in both datasets were used, with negative values of asynchrony (−150, −300, and −500 ms) indicating that the rubber hand was touched first (i.e., the visual signal was presented first), and positive values of asynchrony (+150, +300, and +500 ms) indicating that the participant’s hand was touched first (i.e., the tactile signal was presented first). Each tapping pattern generated in the study consisted of 18 visuo-tactile events occurring over a period of 36 seconds at a 0.5 Hz frequency. This period was substantially extended compared to the 12-second stimulation durations used in the original study (Chancel et al., 2022a) to include a larger number of events, which allowed for greater variance in inter-stimulus intervals—variance necessary for complexity analyses in Simulation 2. Had this extended stimulation been used in the original study, it should not have influenced BCI predictions, given that the BCI model as formulated in Chancel et al. (2022a) is agnostic to the duration of stimulation.

For MCD optimization and fitting in Simulation 1, we generated rhythmic, rectangular visuo-tactile pulses separated by uniform 2 s intervals, designed to match the temporal structure of the visuo-tactile taps used in Chancel et al. (2022a) (Figure 2A). The signal sampling rate was set to 30 Hz, corresponding to the frame rate of visual signal presentation in the original experiment (Chancel et al., 2022a, p. 14). In the original study, tap durations were not measured. Instead, the robotic arms were programmed to stay in the lowest ’tap position’ for 30 ms and leave this position without delay; the actual contact duration may have been somewhat longer, as a slight bend in the probe likely extended the touch beyond the time spent at the lowest position (M. Chancel, personal communication, May 2026). In the face of uncertainty regarding actual tap durations, we used the value closest to the intended stay in the tap position (33 ms), which was the minimal duration that could be represented at a 30 Hz presentation rate. Each visual and tactile tap was therefore represented as a single 33 ms frame, and seven regular tapping patterns were generated, corresponding to the seven asynchrony conditions.

**Figure 2.**
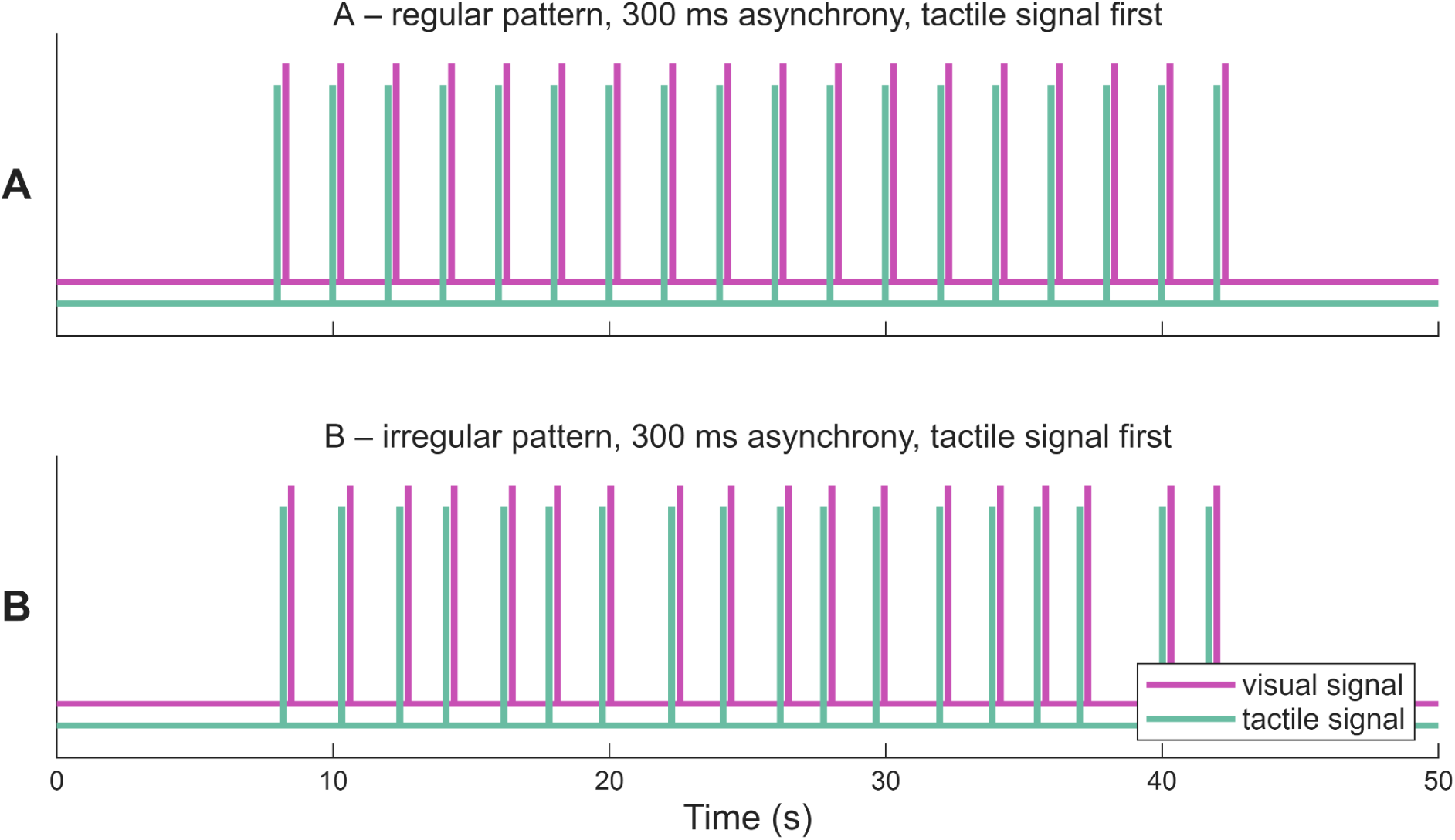
Regular and irregular visuo-tactile patterns used in the study. For Simulation 1, we generated regular visuo-tactile streams simulating tapping patterns from Chancel et al. (2022a). Panel A presents a regular pattern with visual signals (purple dashes) shifted +300 ms with respect to the tactile signals (green dashes). For Simulation 2, 5,000 non-periodic (random and unpredictable) visuo-tactile streams were generated, with the visual signal offset always equal to +300 ms. An example signal is presented in panel B.

The stimulation patterns were padded with a 7-second zero-signal interval at both the beginning and the end to minimize edge artifacts, as recommended by the model’s authors (Parise & Ernst, 2016). All signals were assumed to be free of sensory noise, and the simulated visual signal clearly indicated the presence or absence of a tap.

In Simulation 2, we generated 5,000 irregular visuo-tactile stimulation patterns (Figure 2B). As in Simulation 1, individual taps were modeled as rectangular pulses, and each pattern consisted of 18 taps over 36 seconds, with each tap lasting 33 ms. Unlike the regular stimulation patterns, irregular visuo-tactile streams had inter-stimulus intervals (ISIs) generated stochastically across the entire tapping sequence, without repeating patterns. ISIs across all generated patterns were normally distributed with a mean of 2 seconds and standard deviation of 0.4 seconds. This ensured that approximately 99% of ISI values naturally fell within the [1, 3] second range (mean ± 2.58×SD = [0.97, 3.03]s), with only minimal truncation at the distribution boundaries. Importantly, this ISI distribution was chosen to be consistent with an experimental study that systematically examined the influence of stimulation complexity on RHI strength (Litwin et al., 2020), which used the same stimulation frequency (0.5 Hz) and applied Gaussian jitter to brushstroke onsets, although jitter in Litwin et al. (2020) had a smaller standard deviation (0.2 s).

In each irregular pattern, the visual signal was shifted by a constant delay (300 ms) with respect to the tactile signal. This asynchrony level was selected because it corresponds to the threshold of tolerable visuo-tactile offset for body ownership (Bekrater-Bodmann et al., 2014; Shimada et al., 2009). At this threshold, where causal inference is maximally uncertain, signal complexity could emerge as a relevant factor in determining RHI occurrence, whereas under conditions of perfect synchrony or large temporal mismatch, temporal offset information would dominate. Accordingly, we expected the highest variability in the rate of affirmative responses predicted by the MCD at 300 ms delay, which was required for linear modeling purposes. The tactile signal always occurred first, which was decided arbitrarily.

Zero-signal void periods lasting 7 seconds were embedded both at the beginning and the end of the irregular patterns. The MCD models fitted to each of the two datasets in Simulation 1, with all parameters fixed at their optimal values, were used to predict the probability of experiencing the RHI for each of these irregular patterns and to examine whether the predicted probability was related to the irregularity of the signals.

### Multisensory Correlation Detector Model

The MCD model was implemented following the original computational framework described by Parise and Ernst (2016) (for mathematical formulae related to the MCD implementation, see Appendix 1). Simulated visuo-tactile signals were processed using a single MCD unit, restricting the model to temporal processing and rendering it agnostic to tap locations or spatial noise. Multisensory processing in the MCD begins with independent low-pass filtering of the visual and tactile input signals. These low-pass filters are modeled as exponential functions with modality-dependent time constants, represented by two free parameters of the model: *τ_T_* (time constant for the tactile filter) and *τ_V_* (time constant for the visual filter). Filtered signals are then forwarded to two cross-wired correlation detectors—two subunits which independently combine visuo-tactile information by applying a late-stage low-pass filter to either the visual or the tactile signal. The third free parameter *τ_VT_*corresponds to the time constant of this late-stage filter, which is identical for both subunits. Although the structure of the subunits is mirror-symmetric and shares the same late-stage time constant, differences between *τ_T_* and *τ_V_* lead to modality-specific filtering delays, such that peak integration can occur at small temporal offsets between the signals. Following late-stage filtering, the outputs of the two subunits are multiplied and subtracted, yielding, respectively, time-varying responses that quantify local temporal correlation (*MCD_corr_*) and lag (*MCD_lag_*) between the signals. These responses are averaged over the duration of the analyzed stimulation period to obtain global indices of temporal correlation and temporal offset.

To model RHI detection responses, we assumed that the probability of an affirmative response increases with the temporal correlation between the visual and tactile signals. Within the MCD framework, this correlation is indexed by *MCD_corr_* and signals with highly correlated temporal structures are more likely to be perceived as sharing a common cause (Parise et al., 2012; Parise & Ernst, 2016). However, the MCD architecture also outputs *MCD_lag_*which represents temporal order of the signals (Parise & Ernst, 2016). Even though the *MCD_corr_* seems more directly related to the causal inference problem of determining body ownership, it seems possible that a downstream read-out could use both correlation and lag outputs, especially in the face of visuo-tactile asynchrony. Therefore, we decided to compare two models of RHI detection, which differed in the MCD outputs entering the generalized linear model (GLM) that mapped them onto response probabilities. In the first model (MCD-C), *MCD_corr_* was the only predictor of RHI detection probabilities in the GLM. In the second model (MCD-C+L), RHI detection relies on both correlation and temporal order detection, with both *MCD_corr_* and *MCD_lag_* entering the GLM as predictors (cf. Parise & Ernst, 2025).

### Bayesian Causal Inference model

In the BCI model, the causal inference determining the integration or segregation of visuo-tactile signals depends on the prior probability of a common cause and trial-specific sensory uncertainty. In the original studies on Bayesian modeling of RHI detection probabilities (Chancel & Ehrsson, 2023; Chancel et al., 2022a), BCI models were fitted to all noise conditions. However, because the MCD model could be fitted and evaluated exclusively in the no-noise condition, directly comparing it to original BCI implementations—accounting for all noise conditions but evaluated only in one condition in which MCD could function—would confer a selective advantage for MCD. To ensure a fair comparison, we implemented a BCI model restricted to the no-noise condition (BCI-0), so that it was fitted to and evaluated on the same data as the MCD. BCI-0 has three free parameters: the prior probability of a common cause (*p_same_*), sensory noise (*σ*), and the lapse rate (*λ*). The standard deviation of visuo-tactile asynchronies assumed under separate causes (*σ_s_*) was fixed at 348 ms, the actual standard deviation of the asynchrony levels used in the experiments, as in Chancel et al. (2022a).

### Parameter Estimation

The full MCD modeling pipeline included five (MCD-C) or six (MCD-C+L) free parameters. Three parameters corresponding to the time constants of the filters (*τ_T_*, *τ_V_*, *τ_VT_*) were fitted using a gradient-based constrained optimizer (fmincon) to maximize model fit to response probabilities in the no-noise condition. MCD outputs were transformed into response probabilities using the GLM with the probit link function, including an intercept and either *MCD_corr_* alone (MCD-C) or both *MCD_corr_* and *MCD_lag_* (MCD-C+L) as predictors. MCD outputs were standardized (z-scored) across asynchrony levels before entering the GLM, which improved numerical conditioning without changing the fit or the predictions. The objective function minimized during the optimization was the deviance of the GLM fit, which is equivalent to maximizing the likelihood of the observed responses. Following successful optimization, the same GLM was used to map the MCD outputs, derived from the model with optimized time constants, onto the empirical probabilities of affirmative responses across asynchrony levels. The decisional stage therefore included two (MCD-C) or three (MCD-C+L) additional free parameters corresponding to the linear coefficients (the intercept and the slopes for *MCD_corr_* and, in MCD-C+L, *MCD_lag_*).

For the BCI model, we followed the general fitting procedure described by Chancel et al. (2022a), with two notable exceptions to ensure that the optimization and fitting procedure are fully parallel between MCD and BCI models, rendering the model comparison fair. First, the values of free parameters maximizing the fit of BCI-0 to the observed data were estimated using the same fmincon optimizer as for the MCD, instead of Bayesian Adaptive Direct Search (Acerbi & Ma, 2017) used in the original study. The objective function minimized during optimization was the negative log-likelihood (NLL) of the BCI-0 fit. Second, unlike Chancel et al. (2022a), where parameters were estimated separately for each participant and then averaged, here BCI-0 was fitted to response probabilities aggregated across participants at each asynchrony level, exactly as the MCD models.

The optimization procedure was identical for the MCD and BCI models, and each model was fitted separately to each dataset. Parameters were estimated with fmincon, which searches for the minimum of the objective function by following its local gradient within hard bounds on parameter values. Both objective functions were deterministic, smooth, and fast to evaluate (about 4 ms per evaluation for the MCD and 0.1 ms for BCI-0), which created a setting in which gradient-based optimization is efficient and reliable. Because a gradient-based optimizer converges to the minimum nearest to its starting point, the optimization procedure was repeated 100 times, with each run initialized with a different parameter combination. The solution yielding the minimal objective value was retained. The robustness of the optimization procedure was assessed using the number of times in which the algorithm converged to the same optimal parameter set by reaching an objective function value within 0.001 of the minimum across all runs.

Initial parameter values were always drawn uniformly from within the plausible bounds for the relevant parameters, whereas hard bounds defined the limits of the parameter space explored by the optimizer. For MCD, parameter bounds were approximated based on prior modeling work (Parise & Ernst, 2016; Pesnot Lerousseau et al., 2022), whereas for BCI both hard and plausible bounds were set identically to those used in Chancel et al. (2022a). Bounds used in the optimization are presented in Table 1.

**Table 1.**
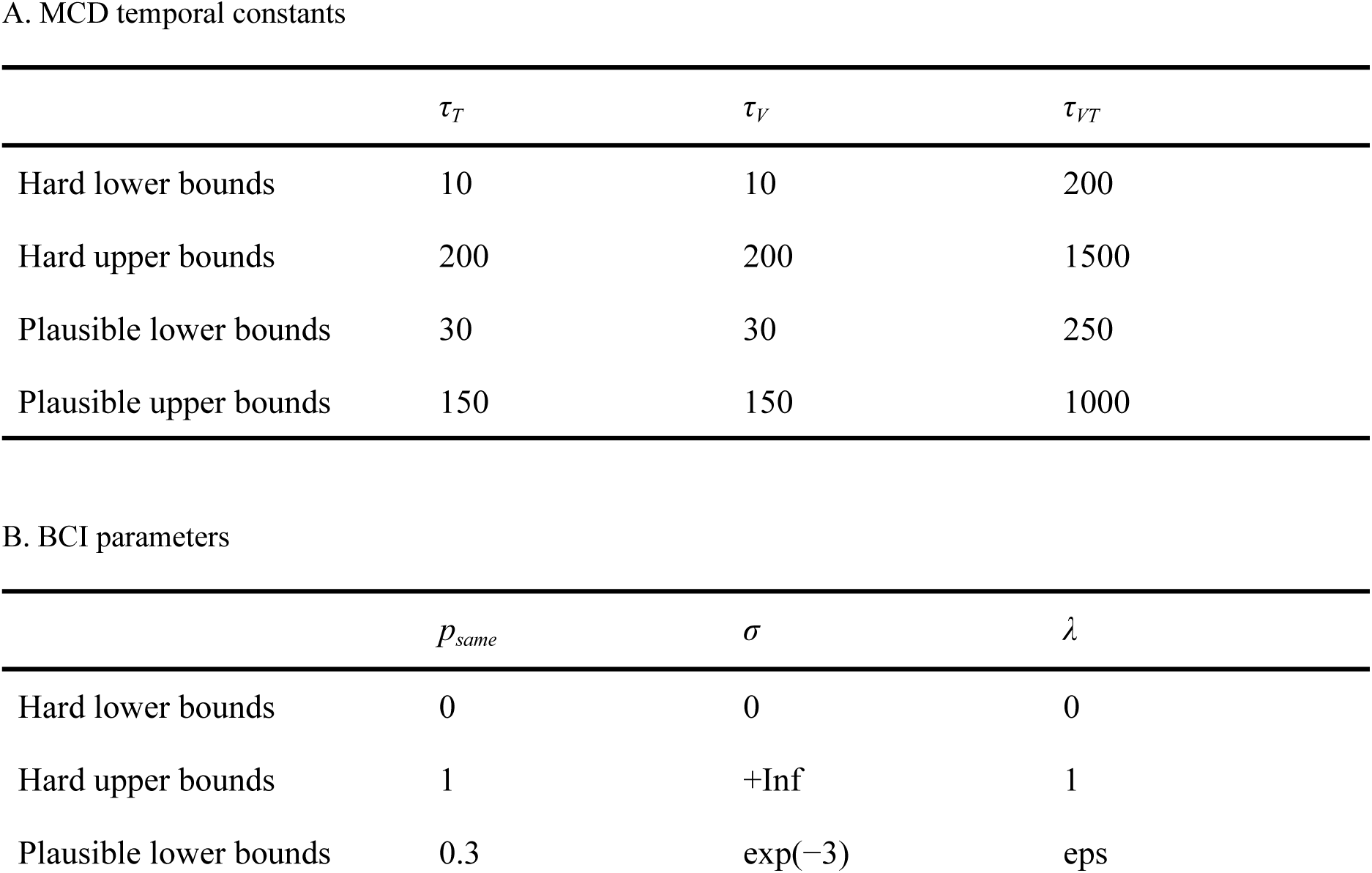
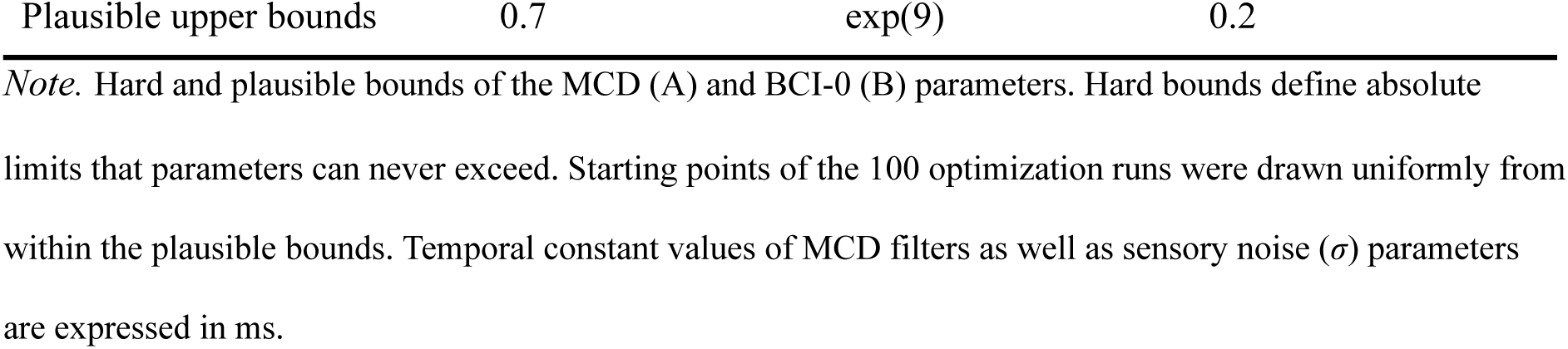
Hard and plausible bounds of model parameters.

| A. MCD temporal constants |  |  |  |
|  | *τ<sub>T</sub>* | *τ<sub>V</sub>* | *τ<sub>VT</sub>* |
|---|---|---|---|
| Hard lower bounds | 10 | 10 | 200 |
| Hard upper bounds | 200 | 200 | 1500 |
| Plausible lower bounds | 30 | 30 | 250 |
| Plausible upper bounds | 150 | 150 | 1000 |
| B. BCI parameters |  |  |  |
|  | *p<sub>same</sub>* | *σ* | *λ* |
| Hard lower bounds | 0 | 0 | 0 |
| Hard upper bounds | 1 | +Inf | 1 |
| Plausible lower bounds | 0.3 | exp(−3) | eps |
| Plausible upper bounds | 0.7 | exp(9) | 0.2 |

*Note.* Hard and plausible bounds of the MCD (A) and BCI-0 (B) parameters. Hard bounds define absolute limits that parameters can never exceed. Starting points of the 100 optimization runs were drawn uniformly from within the plausible bounds. Temporal constant values of MCD filters as well as sensory noise (*σ*) parameters are expressed in ms.

### Model Comparison and Cross-Validation

Simulation 1 served two purposes: to compare the MCD with BCI-0 in fitting and predicting RHI detection probabilities, and to select the MCD model used in Simulations 2 and 3 based on their in- and out-of-sample performances. Model fits were compared using multiple fit metrics, including the NLL of the observed responses, the Akaike Information Criterion (AIC; Akaike, 1973), and the Bayesian Information Criterion (BIC; Schwarz, 1978), which additionally penalizes model complexity as reflected in the number of free parameters. As a measure of absolute model performance, we computed Nagelkerke’s pseudo-R² (Nagelkerke, 1991) relative to an intercept-only model that predicts the overall proportion of affirmative responses at every asynchrony level.

To assess how well the models predict RHI detection at asynchrony levels they were not fitted to, we used leave-one-asynchrony-out cross-validation within each dataset. In each of seven folds, one asynchrony level was omitted, and the complete fitting procedure (100 optimization runs, with time constants and GLM coefficients refitted for the MCD) was repeated on the remaining six levels. Each fitted model then predicted the RHI probability at the omitted level. When repeated for all asynchrony levels, this procedure yielded a psychometric curve in which every point was an out-of-sample prediction, testing whether a model can reproduce the shape of the observed psychometric curve—including its asymmetry—from the remaining asynchrony levels only. Out-of-sample performance was quantified with the NLL of these predictions and with Nagelkerke’s pseudo-R², with the intercept-only model cross-validated in the same way (i.e., estimated on the six training levels of each fold).

### Signal Complexity

To quantify signal complexity for each simulated visuo-tactile pattern in Simulation 2, we followed a method originally developed for audiovisual streams (Denison et al., 2013).

Signal complexity was operationalized using the information-theoretic entropy measure based on ISI durations. Two entropy measures were computed. First-order ISI entropy was calculated using the following formula:

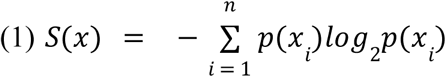

where *n* denotes the number of unique ISIs in the visuo-tactile stream, each *x_i_* represents a unique ISI duration, and *p*(*x_i_*) stands for the probability of each unique ISI occurrence. First-order ISI entropy therefore captures signal complexity in terms of variability in ISI durations; however, identical sets of ISIs with the same probability distribution can give rise either to rhythmic patterns (with all ISIs repeatedly reappearing in the same order) or non-rhythmic, temporally unpredictable patterns. To account for this unpredictability, we calculated a second-order ISI entropy that reflected variability in pairs of consecutive ISI durations (first–second, then second–third, etc.). Second-order entropy was calculated using the same formula, but with each *x_i_* standing for one of the unique ISI pairs present in the stream.

Using ISI values at full temporal resolution (1/30 s precision) would result in most ISIs being unique within every generated signal, yielding maximal entropy values in nearly all signals. This would prevent meaningful discrimination among the generated visuo-tactile signals in terms of complexity. Therefore, to ensure actual variability in signal complexity across irregular patterns, ISI values were rounded. To balance discriminability of patterns against information loss due to rounding, ISIs were rounded to the nearest 0.2 s for first-order entropy (yielding 11 possible ISI values between 1 and 3 s) and to the nearest 0.5 s for second-order entropy (yielding 5 possible values per ISI and thus 25 possible ISI pair combinations). A coarser bin width was used for second-order entropy because the number of possible ISI pairs grows with the square of the number of possible ISI values; with 0.2 s bins, most of the 16 consecutive ISI pairs in each pattern would be unique, which would reduce the variability of second-order entropy across patterns. For regular patterns, in which visuo-tactile signals repeated every 2 seconds, there was no variance in either ISI durations or ISI pairs, and both first-order and second-order entropy were equal to 0.

Using 5,000 irregular stimulation patterns as data points, we first compared MCD-predicted RHI probabilities for irregular patterns with the prediction for the regular pattern with the same asynchrony, using a one-sample t-test with Cohen’s d as the effect size. We then examined the relationship between signal complexity and MCD-predicted RHI occurrence probability using multiple linear regression. The dependent variable was a continuous probability estimate bounded between 0 and 1. The regression model included an intercept and first- and second-order ISI entropy as predictors, without an interaction term.

The linear regression fit was assessed using adjusted R^2^, which quantifies the proportion of explained variance. Semi-partial correlations were calculated to determine the unique variance explained by each entropy measure independently of the other. To test whether the relationship originated in the MCD unit rather than in the GLM read-out, we also computed Pearson correlations between each entropy measure and the raw *MCD_corr_* output. All analyses were performed separately for the MCD-C model fitted to each dataset.

### Point of Strongest Ownership and Windows of Subjective Ownership

Parameters *τ_V_* and *τ_T_* characterize the time-dependent behavior of low-pass filters, with lower time constant values related to quicker stimulus processing (Smith, 1997). Therefore, varying time constant parameters should systematically alter effective processing times of visual and tactile signals, and, in turn, MCD-predicted RHI detection probabilities. To examine this hypothesis, we calculated psychophysical RHI measures related to the *point of strongest ownership* (PSO) and *window of subjective ownership* (WSO). These measures were modeled after standard psychophysical measures used in simultaneity judgment tasks, but ownership-specific terms were introduced to reflect the distinction between visuo-tactile simultaneity perception and visuo-tactile integration underlying body ownership. PSO was defined as visuo-tactile delay at which the probability of RHI detection was highest, with values deviating from 0 indicating that the MCD predicts a non-zero temporal offset between signals for the highest RHI probability. WSO was defined as the temporal interval during which stimuli from different senses can be integrated to a sufficient extent to reliably elicit affirmative responses in a RHI detection task, analogous to how visuo-tactile temporal binding windows reflect intervals of consistently perceived simultaneity. Based on the relation between time constant values and stimulus processing times, we expected that (1) large time constant asymmetries should produce skewed RHI probabilities, with PSO values shifting toward earlier presentation of the modality with higher time constants to compensate for longer processing times, and (2) generally lower time constants, indicating faster unisensory processing, should produce narrower WSOs.

To examine these relationships, we manipulated the time constants of the MCD model in two separate simulations. For each dataset, the optimal MCD-C parameter values found in Simulation 1 were used as the baseline parameter set. For PSO analyses, we generated eight parameter sets by progressively subtracting or adding 15 ms to the baseline value of the tactile filter time constant, while keeping the visual filter time constant unchanged. For WSO analyses, we generated eight additional parameter sets: four with progressively lower time-constant values and four with progressively higher values. These values were obtained by successively subtracting or adding 15 ms to the baseline parameter values of both visual and tactile filters jointly. In both simulations, the late-stage filter time constant (*τ_VT_*) was held fixed at its baseline value.

Each generated parameter set was used to define a separate MCD model, totaling 16 additional MCD models per dataset for PSO and WSO analyses. Using each of these MCD models, we (1) estimated visuo-tactile correlations (*MCD_corr_*) across the seven visuo-tactile asynchrony levels using the rhythmic visuo-tactile pattern generated in Simulation 1, (2) fitted a GLM with a probit link function to relate these *MCD_corr_* values to the response probabilities of the respective dataset, thereby estimating the linear model parameters (intercept and *MCD_corr_* slope) for each modified MCD, and (3) generated predictions of RHI occurrence probabilities across all asynchrony levels. For each MCD model, predicted RHI probabilities were subsequently modeled as a function of asynchrony using asymmetric psychometric functions defined as the difference of two cumulative Gaussian functions (Parise & Ernst, 2025):

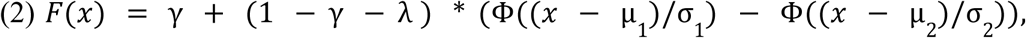

where *Φ* is the cumulative normal distribution function, *μ_1_* and *μ_2_* are the means of the two cumulative Gaussians, *σ_1_*and *σ_2_* are their standard deviations, *γ* is the guess rate (lower asymptote), and *λ* adjusts the upper asymptote. This function allowed for asymmetry in the psychometric curve and enabled extraction of PSO and WSO values for each MCD model.

PSO was operationally defined as the asynchrony value at which the predicted RHI detection probability reached its peak, whereas WSO was defined as the half-width of the fitted function at half of its maximum height above the lower asymptote (cf. Parise & Ernst, 2025). Psychometric functions were fitted to the predicted probabilities by least squares from 20 starting points, and the best-fitting solution was retained.

All simulations and analyses were performed using MATLAB R2024a (The MathWorks, Inc., Natick, MA, USA). Claude Opus 5.5 or earlier versions (Anthropic, San Francisco, CA, USA) were used to generate parts of the MATLAB code and for AI-assisted copy editing.

## Results

### Simulation 1

The goal of Simulation 1 was to compare the relative fits of the MCD models and BCI-0 to empirical RHI detection probabilities, separately for each dataset. For MCD-C, the same optimal parameter set was reached in 97 of 100 optimization runs for the first dataset (*τ_T_* = 121 ms, *τ_V_* = 128 ms, *τ_VT_* = 496 ms) and in 64 runs for the second dataset (*τ_T_* = 123 ms, *τ_V_* = 105 ms, *τ_VT_* = 408 ms). For MCD-C+L, the minimum was reached in 94 and 90 runs, with time constants similar to those of MCD-C (first dataset: *τ_T_* = 121 ms, *τ_V_* = 125 ms, *τ_VT_* = 482 ms; second dataset: *τ_T_* = 126 ms, *τ_V_* = 103 ms, *τ_VT_* = 433 ms). All runs that reached the minimum converged to the same parameter values.

For BCI-0, the minimum NLL was reached in 87 runs for the first dataset (*p_same_* = 0.70, *σ* = 130 ms, *λ* = 0.14) and in 22 runs for the second dataset (*p_same_* = 0.48, *σ* = 144 ms, *λ* = 0.21). For the first dataset, the estimates indicate a prior belief that visual and tactile signals were more likely to originate from a common source and a relatively low lapse rate, in line with Chancel et al. (2022a). In the second dataset, the prior probability of a common cause was lower, indicating similar probabilities of common and separate causes, also consistent with the original study which found relatively low *p_same_* = .55 in the no-noise condition (Chancel & Ehrsson, 2023, p. 76). These findings indicate that our optimization procedure produced results comparable to those of the original studies, even though they used a different algorithm and estimated parameters separately for each participant, only averaging them at the end.

In both datasets, the MCD models fitted RHI detection probabilities more closely than BCI-0, producing near-perfect fits (Figure 3). In the first dataset, MCD-C reached a lower NLL (17.90 vs. 21.40), a higher pseudo-R² (0.948 vs. 0.937), and a lower AIC (45.80 vs. 48.80) than BCI-0, whereas its larger number of free parameters resulted in a higher BIC (71.49 vs. 64.22). Adding *MCD_lag_* to the GLM barely improved the fit (ΔNLL = 0.05), resulting in higher AIC (47.70) and BIC (78.53) values for MCD-C+L compared to MCD-C due to the higher number of parameters. In the second dataset, MCD-C outperformed BCI-0 on all metrics, by a substantially larger margin (NLL: 24.24 vs. 47.75; pseudo-R²: 0.957 vs. 0.914; AIC: 58.47 vs. 101.50; BIC: 89.23 vs. 119.95). Again, even though introducing *MCD_lag_*produced a small improvement in fit (ΔNLL = 0.13), it failed to compensate for the loss of parsimony as evidenced by information criterion metrics (AIC: 60.21; BIC: 97.13). Both MCD models reproduced the asymmetry of the observed psychometric curves, including its reversal between datasets: RHI probabilities were higher when the visual signal led in the first dataset and when the tactile signal led in the second dataset. BCI-0, whose predictions were symmetric around zero asynchrony by construction, could not capture either asymmetry.

**Figure 3.**
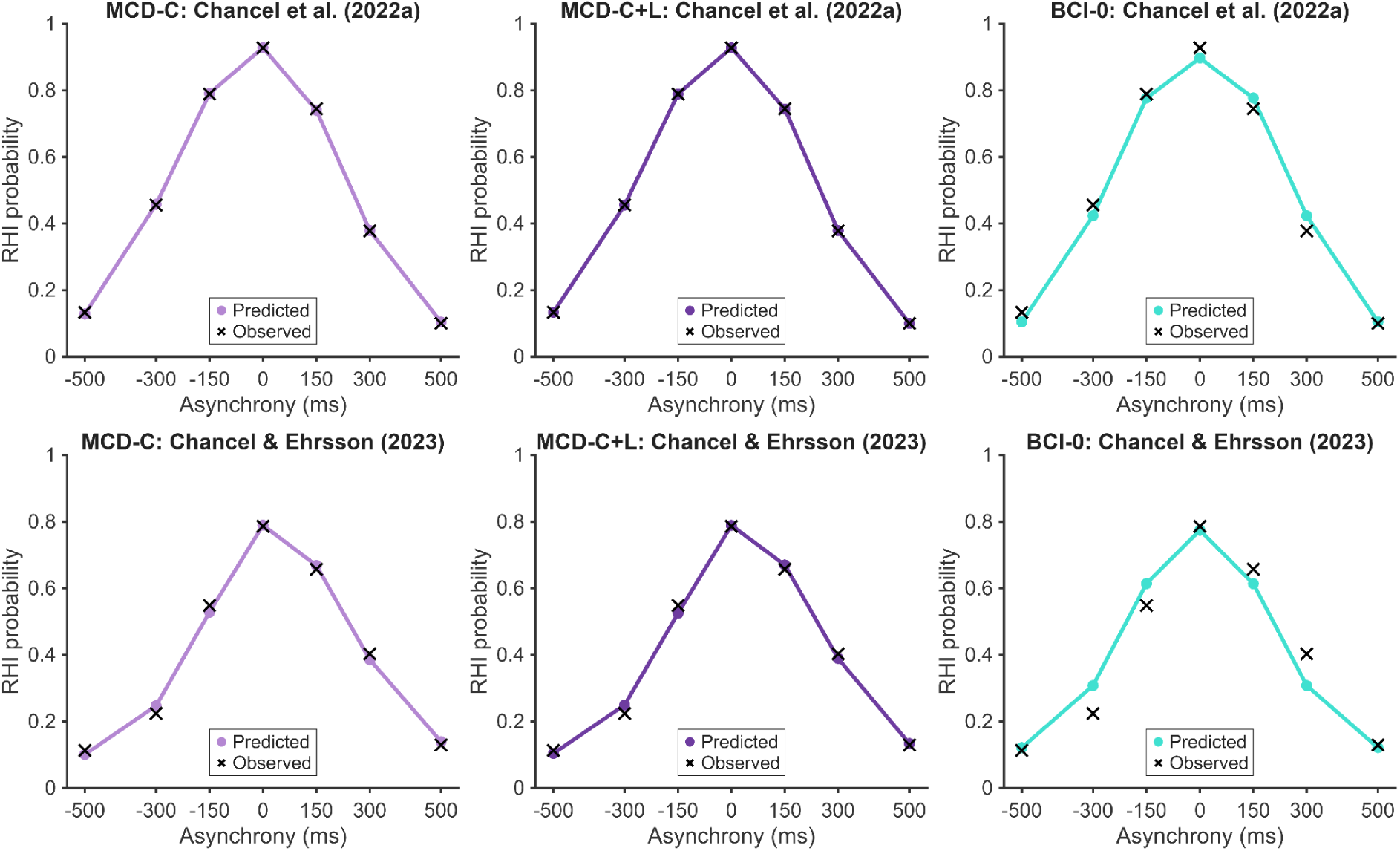
Fits of the MCD and BCI-0 models to RHI detection probabilities in both datasets. Rows: Chancel et al. (2022a) and Chancel and Ehrsson (2023); columns: MCD-C (light purple), MCD-C+L (dark purple), and BCI-0 (teal). Colored dots and lines represent model predictions after fitting to all seven asynchrony levels; black crosses represent observed RHI probabilities. Observed probabilities are proportions of affirmative responses aggregated across participants (Chancel et al., 2022a: 15 participants, 180 trials per asynchrony level; Chancel and Ehrsson, 2023: 31 participants, 496 trials per asynchrony level).

**Figure 4.**
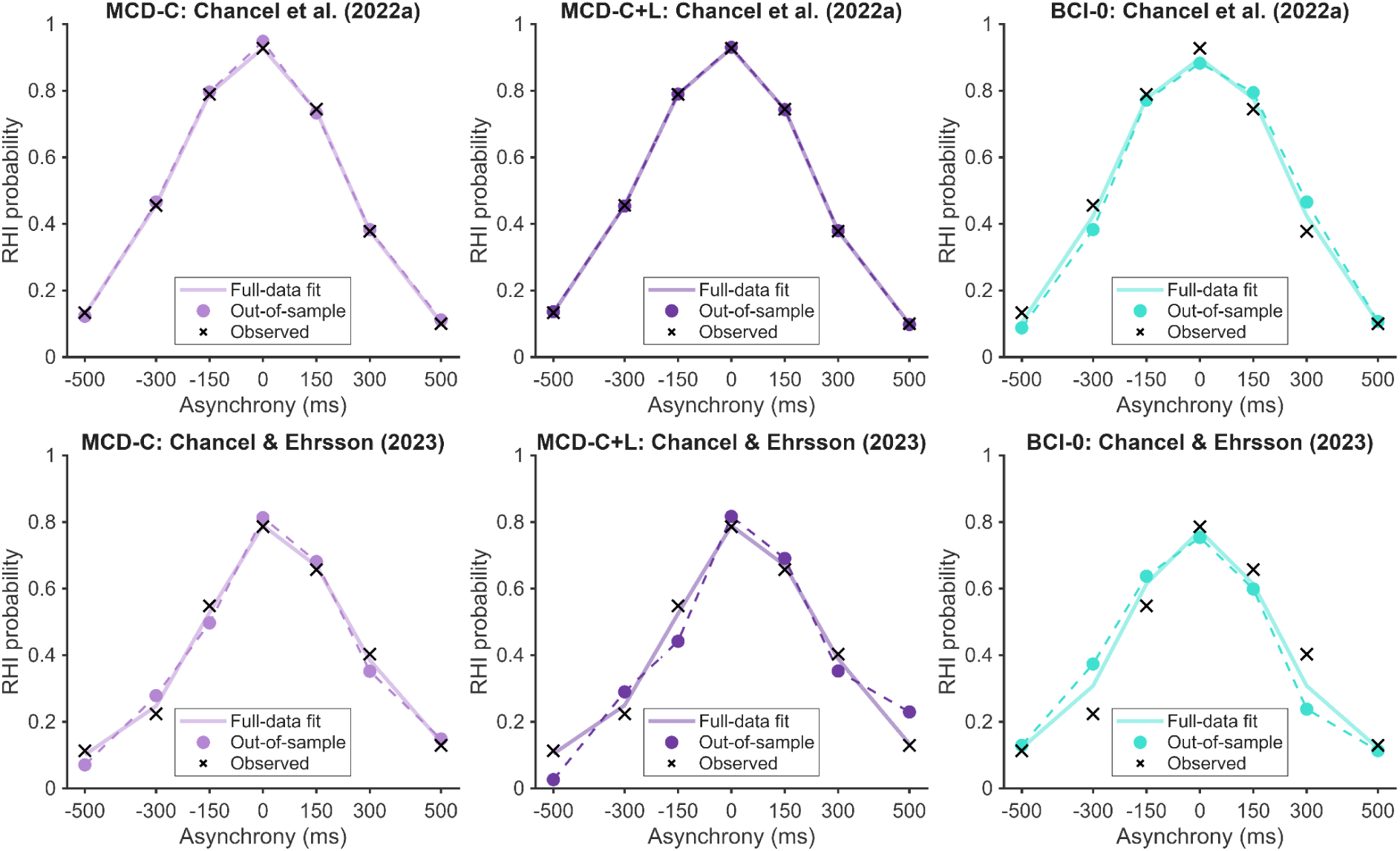
Out-of-sample predictions of RHI detection probabilities in leave-one-asynchrony-out cross-validation. Rows: Chancel et al. (2022a) and Chancel and Ehrsson (2023); columns: MCD-C, MCD-C+L, and BCI-0. Filled dots connected by dashed lines are out-of-sample predictions, each obtained from a model fitted to the remaining six asynchrony levels. Light solid lines show the fits to all seven levels (Figure 3) for reference. Black crosses represent observed RHI probabilities. MCD-C predicts RHI detection probabilities at omitted asynchrony levels, including the direction of the asymmetry, better than BCI-0 in both datasets.

Another goal of Simulation 1 was to validate models by examining their predictions at asynchrony levels omitted from fitting, again using both datasets. In leave-one-asynchrony-out cross-validation, MCD-C predicted the omitted asynchrony levels better than BCI-0 in both datasets (first dataset: NLL = 18.87 vs. 28.22, pseudo-R² = 0.961 vs. 0.941; second dataset: NLL = 39.58 vs. 95.27, pseudo-R² = 0.949 vs. 0.875). As in the fitting procedure spanning all asynchronies, the out-of-sample predictions of MCD-C preserved the direction of the asymmetry in both datasets. In contrast, BCI-0 tended to mispredict the omitted levels in the direction opposite to the observed asymmetry in both datasets, with the largest errors around the inflection points of the RHI probability distribution (±300 ms levels). MCD-C+L generalized inconsistently: it performed slightly better than MCD-C in the first dataset (NLL = 17.87, pseudo-R² = 0.963), but worst of all models in the second dataset (NLL = 101.42, pseudo-R² = 0.867). Because adding *MCD_lag_* neither improved the fit nor generalized reliably, MCD-C was used in Simulations 2 and 3. The results of the fitting procedure as captured by all fit metrics in both fitting and leave-one-out cross-validation procedures are summarized in Table 2.

**Table 2.** Fits to all asynchrony levels and leave-one-asynchrony-out cross-validation.

| Dataset | Model | k | NLL | AIC | BIC | Pseudo-R² | CV NLL | CV pseudo-R² |
|---|---|---|---|---|---|---|---|---|
| Chancel et al. (2022a) | MCD-C | 5 | 17.90 | 45.80 | 71.49 | 0.948 | 18.87 | 0.961 |
|  | MCD-C+L | 6 | 17.85 | 47.70 | 78.53 | 0.948 | 17.87 | 0.963 |
|  | BCI-0 | 3 | 21.40 | 48.80 | 64.22 | 0.937 | 28.22 | 0.941 |
| Chancel & Ehrsson (2023) | MCD-C | 5 | 24.24 | 58.47 | 89.23 | 0.957 | 39.58 | 0.949 |
|  | MCD-C+L | 6 | 24.11 | 60.21 | 97.13 | 0.957 | 101.42 | 0.867 |
|  | BCI-0 | 3 | 47.75 | 101.50 | 119.95 | 0.914 | 95.27 | 0.875 |

*Note.* k: number of free parameters; NLL: negative log-likelihood; AIC: Akaike Information Criterion; BIC: Bayesian Information Criterion; pseudo-R²: Nagelkerke’s pseudo-R²; CV: cross-validation. Cross-validated pseudo-R² was computed relative to an intercept-only model cross-validated in the same way and is therefore not directly comparable with the in-sample pseudo-R².

### Simulation 2

The aim of Simulation 2 was to determine whether MCD predictions of RHI probability vary with the complexity of the visuo-tactile signals used to elicit the RHI. To this end, we applied two MCD-C models fitted to both datasets in Simulation 1, with all parameters fixed, to 5,000 irregular visuo-tactile tapping patterns with a fixed 300 ms tactile-first asynchrony, and predicted the proportion of ‘yes’ responses for each pattern. In both cases, the mean predicted RHI probability across irregular patterns was higher than the probability predicted for the regular pattern with the same asynchrony, with very strong effect sizes (first dataset: *M* = 0.495, *SD* = 0.066, vs. 0.379; *t*(4999) = 123.43, *p* < 0.001, *d* = 1.75; second dataset: *M* = 0.481, *SD* = 0.052, vs. 0.386; *t*(4999) = 129.38, *p* < 0.001, *d* = 1.83; Figure 5). Notably, the BCI models would predict the same proportion of RHI detection responses for each of the 5,000 irregular patterns, given that the asynchrony was equal in all of them.

**Figure 5.**
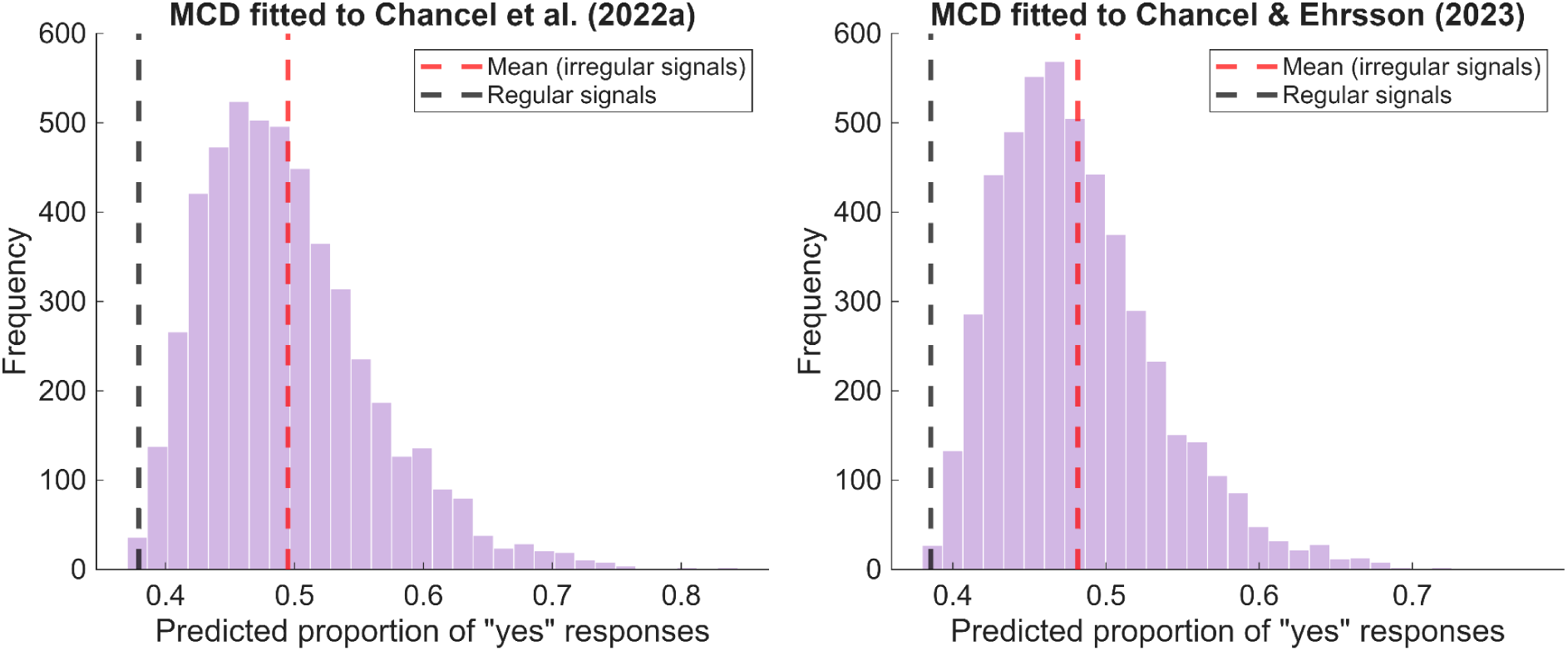
RHI probabilities predicted by the MCD-C models for 5,000 irregular visuo-tactile patterns. Left: MCD-C fitted to Chancel et al. (2022a); right: MCD-C fitted to Chancel and Ehrsson (2023). The red dashed line represents the mean predicted RHI probability for 5,000 irregular visuo-tactile patterns; the black dashed line shows the prediction for the regular pattern with the same asynchrony (+300 ms, tactile signal first). Histograms show the distribution of predicted probabilities across the irregular patterns (*N* = 5,000 per panel). MCD-C predicts higher RHI probabilities for irregular visuo-tactile signals compared to the rhythmic ones. The mean prediction for irregular patterns was compared with the prediction for the regular pattern using one-sample t-tests (first dataset: *t*(4999) = 123.43, *p* < 0.001, *d* = 1.75; second dataset: *t*(4999) = 129.38, *p* < 0.001, *d* = 1.83).

To further examine the relationship between signal complexity and MCD-predicted RHI probability, we calculated two information-theoretic measures of temporal signal complexity for each generated tapping pattern. First-order entropy reflected the degree of irregularity of signals (defined as variability in ISIs), and second-order entropy reflected the degree of unpredictability (defined as variability in ISI pairs). The generated visuo-tactile streams exhibited varying levels of both irregularity (mean first-order ISI entropy = 2.63 bits; SD = 0.24) and unpredictability (mean second-order ISI entropy = 2.88 bits; SD = 0.36).

Having established first- and second-order entropies for all irregular signals, we fitted a multiple linear regression model with both signal complexity measures entered as predictors and MCD-predicted RHI probability as the dependent variable. No interaction term was included in the model.

For the MCD-C model fitted to the first dataset, both first-order entropy (*b* = 0.077, *SE* = 0.004; *t*(4997) = 17.24; *p* < 0.001) and second-order entropy (*b* = 0.045, *SE* = 0.003; *t*(4997) = 15.20; *p* < 0.001) were significant predictors of RHI detection probability, even when accounting for each other within the same model. The model explained 22.7% of the variance in MCD-predicted RHI probabilities, and the proportions of variance uniquely explained by first-order entropy (*sr*² = 0.046) and second-order entropy (*sr*² = 0.036) were comparable. The MCD-C model fitted to the second dataset yielded the same pattern: first-order entropy (*b* = 0.066, *SE* = 0.003; *t*(4997) = 19.20; *p* < 0.001) and second-order entropy (*b* = 0.039, *SE* = 0.002; *t*(4997) = 16.86; *p* < 0.001) were both significant predictors, the model explained 26.6% of the variance, and the two entropy measures again explained comparable amounts of unique variance (first-order entropy: *sr*² = 0.054; second-order entropy: *sr*² = 0.042). Both entropy measures already correlated with the raw *MCD_corr_* output to a similar degree (first dataset: *r*(4998) = 0.435 and 0.423; second dataset: *r*(4998) = 0.473 and 0.460; all *p* < 0.001), indicating that the effect originated in the MCD unit rather than in the GLM read-out (Figure 6). Taken together, these findings indicate that, for both parameter sets, the MCD predicts both irregularity and unpredictability to foster integration of visuo-tactile signals for RHI, each making a unique contribution.

**Figure 6.**
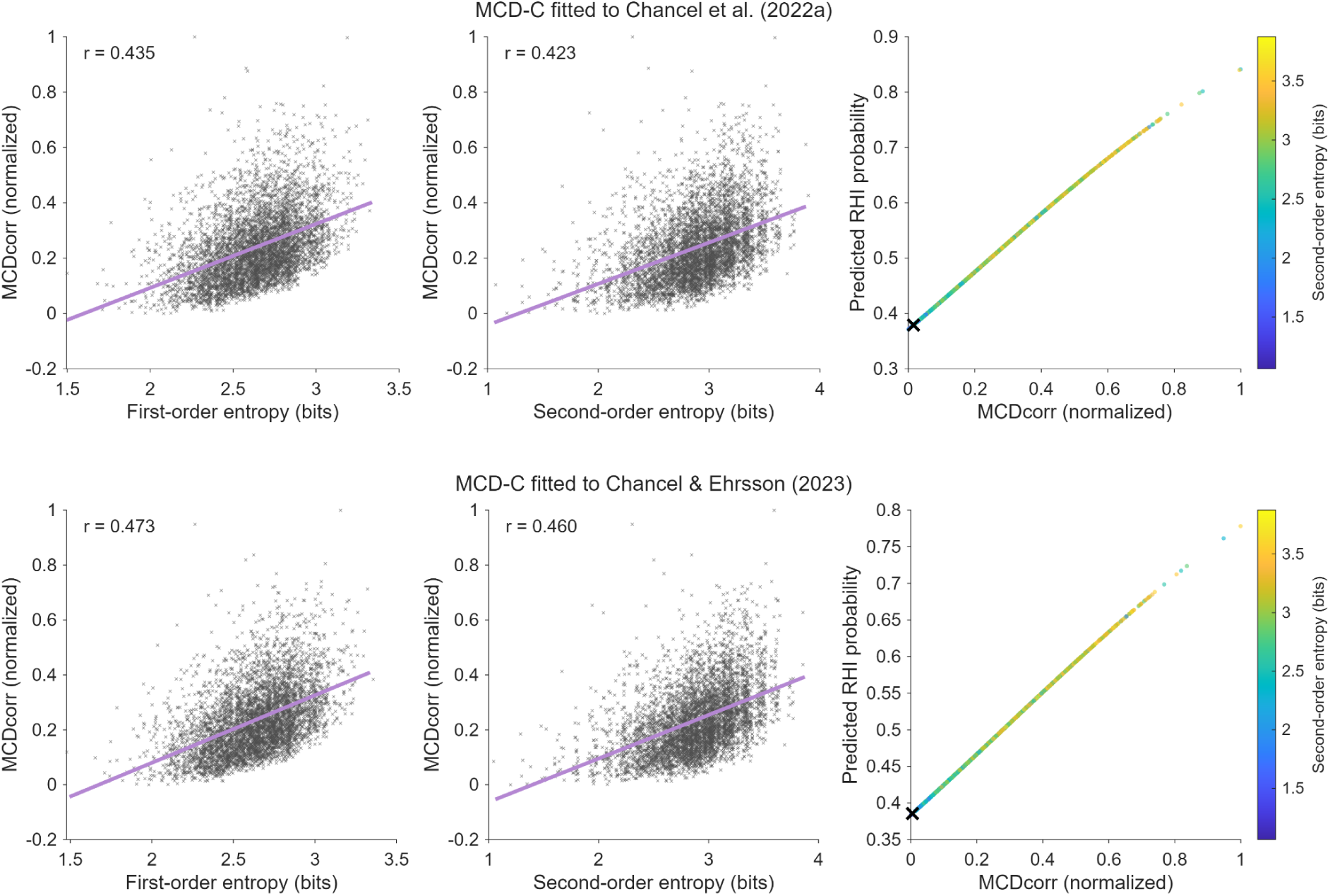
Signal complexity, *MCD_corr_*, and predicted RHI probability. Rows: MCD-C fitted to Chancel et al. (2022a) and to Chancel and Ehrsson (2023). Left and middle columns: *MCD_corr_* output for 5,000 irregular patterns as a function of first- and second-order ISI entropy, with least-squares lines and Pearson correlations (*N* = 5,000 patterns, all *p* < 0.001); entropy values are jittered horizontally for visibility. Right column: RHI probability predicted by the GLM read-out as a function of *MCD_corr_*, colored by second-order entropy; the black cross marks the regular pattern with the same asynchrony. *MCD_corr_* is min–max normalized across the 5,000 irregular patterns of each parameter set (0 = lowest, 1 = highest value), because its raw scale depends on the time constants, number of events, and signal duration. More complex visuo-tactile patterns produce higher *MCD_corr_*, which the GLM read-out translates into higher predicted RHI probability.

### Simulation 3

In Simulation 3, we examined whether varying the time constants of the MCD-C model systematically changes predicted RHI probabilities across visuo-tactile asynchrony conditions, starting from the parameters fitted to each dataset in Simulation 1. First, we manipulated only the tactile filter time constant, in 15 ms steps up to ±60 ms around its baseline value. Predicted PSO values changed monotonically with *τ_T_* in both datasets (Figure 7; Figures S1 and S2), from −79 to 72 ms for the first dataset and from −66 to 122 ms for the second dataset. Shorter tactile time constants shifted the PSO toward visual-first asynchronies, requiring increasingly earlier visual signals to achieve the highest RHI probability. On the other hand, longer tactile time constants shifted the PSO toward tactile-first asynchronies, requiring increasingly earlier tactile signals to achieve the highest RHI probability. Larger time constant asymmetries produced more skewed psychometric functions and stronger PSO shifts toward the sensory modality with the higher time constant, compensating for its slower processing. The baseline parameter sets follow the same relation: in the first dataset, *τ_T_* was lower than *τ_V_* (121 vs. 128 ms) and the PSO was negative (−11 ms), whereas in the second dataset, *τ_T_* was higher than *τ_V_* (123 vs. 105 ms) and the PSO was positive (31 ms). Through the lens of MCD, the opposite asymmetries observed in the two datasets thus correspond to opposite relations between the tactile and visual time constants, and *MCD_lag_*output—encoding temporal order of the signals—is not required for these asymmetries to be captured.

**Figure 7.**
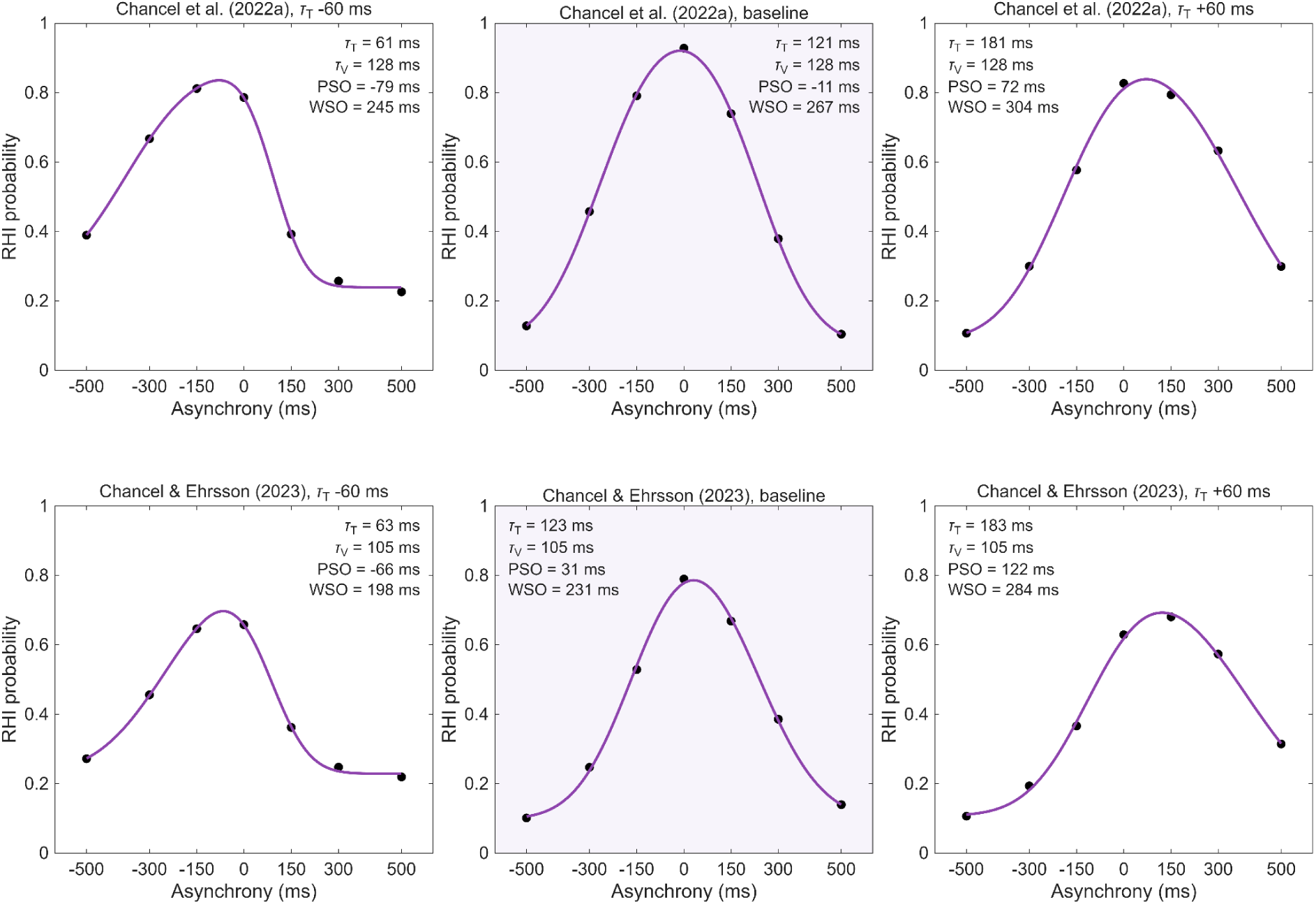
Shifting PSOs by varying the time constant of the tactile filter. Rows: MCD-C fitted to Chancel et al. (2022a) and to Chancel and Ehrsson (2023). Columns: tactile time constant decreased by 60 ms, baseline parameter set from Simulation 1 (shaded), and tactile time constant increased by 60 ms; the visual and late-stage time constants were kept at their baseline values. Black dots represent MCD-predicted RHI probabilities and purple lines the fitted asymmetric psychometric functions. Values in each panel give the early filter time constants and the resulting PSO and WSO. All nine parameter sets for each dataset are shown in Figures S1 and S2.

Next, we manipulated the time constants of the tactile and visual filters jointly (Figure 8). Joint manipulation did not considerably change PSO values, which remained slightly negative for the first dataset (−16 to −11 ms) and positive for the second dataset (12 to 32 ms). In contrast, WSO values increased with the time constants in both datasets (first dataset: 185–308 ms; second dataset: 139–287 ms; Figure 8; Figures S3 and S4). These results suggest that the MCD model predicts higher time constant values for the low-pass filters, indicating slower unisensory processing, to result in wider windows of visuo-tactile integration for body ownership. Importantly, this interpretation is consistent with increased WSO values observed after increasing the tactile filter time constant only (Figure 7), which also contributed to overall longer unisensory processing.

**Figure 8.**
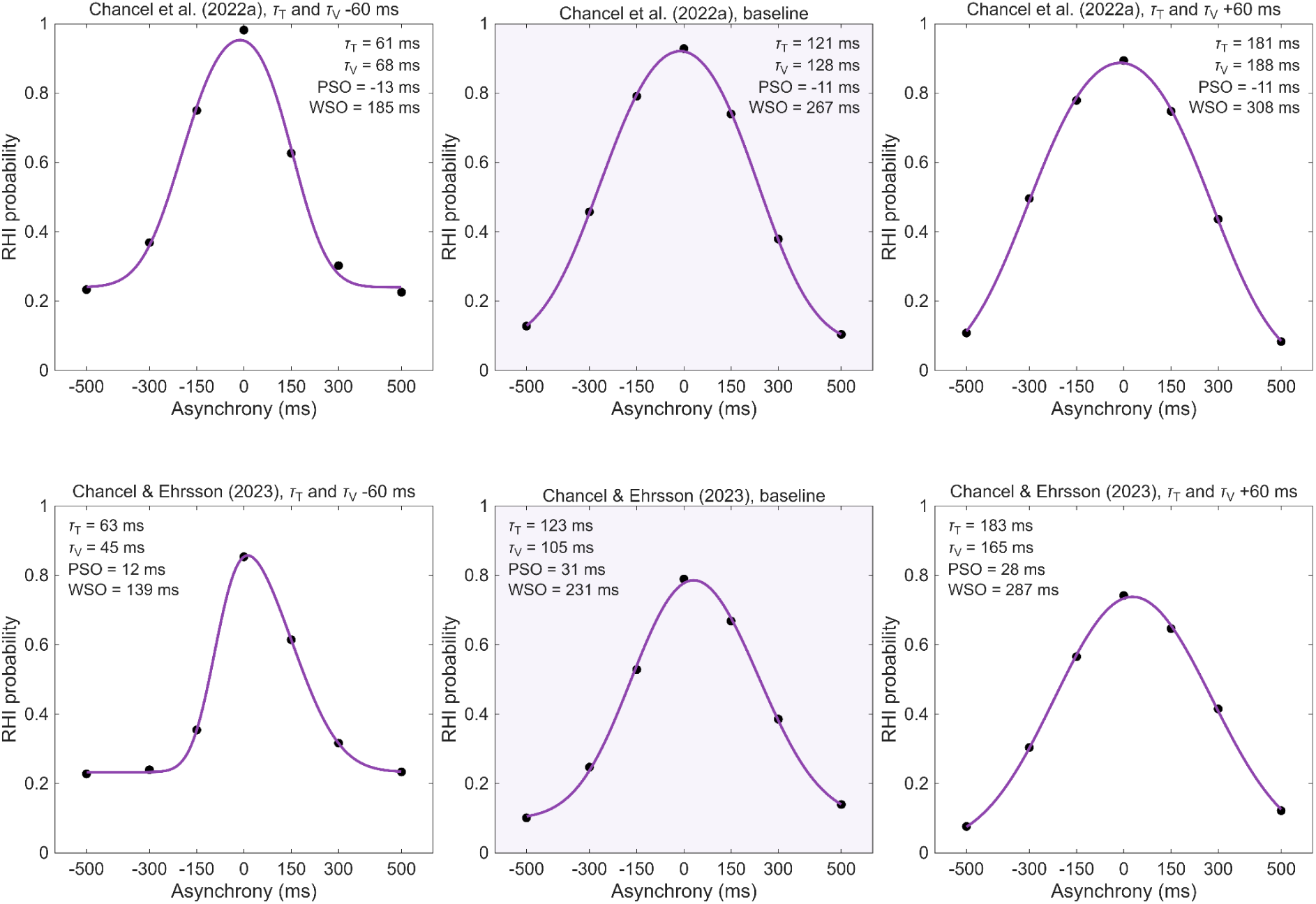
Shifting WSOs by varying the time constants of both early filters jointly. Rows: MCD-C fitted to Chancel et al. (2022a) and to Chancel and Ehrsson (2023). Columns: tactile and visual time constants jointly decreased by 60 ms, baseline parameter set from Simulation 1 (shaded), and both time constants jointly increased by 60 ms; the late-stage time constant was kept at its baseline value. Lower time constants produced narrower WSOs, meaning that signals must be presented closer in time to reliably elicit the RHI. All nine parameter sets for each dataset are shown in Figures S3 and S4.

## Discussion

The present study investigated the applicability of the MCD model (Parise & Ernst, 2016) for explaining response probabilities in the RHI detection task. The results demonstrated that the MCD provided excellent fits to psychophysical RHI detection data from two independent datasets and performed well in leave-one-asynchrony-out cross-validation, successfully predicting RHI detection probabilities at omitted asynchrony levels. Additionally, MCD predicted visuo-tactile tapping patterns with complex temporal structures to yield higher rates of RHI occurrence compared to rhythmic tapping patterns. We also demonstrated that manipulating the temporal constants of visual and tactile filters in the MCD unit systematically affected RHI susceptibility under different asynchrony conditions, accounting for possible asymmetries of RHI probability distribution. Together, these results indicate that the MCD model offers a versatile computational account of temporal visuo-tactile integration underlying body ownership.

Three computational simulations were carried out in this study. In Simulation 1, we compared two variants of the MCD model (MCD-C and MCD-C+L) and the BCI-0 model fit to the no-noise condition only with respect to several fit metrics, using two independent RHI detection datasets. In the first dataset, MCD-C performed better when assessed using pseudo-R² or AIC; however, it incurred a stronger complexity penalty under BIC compared to BCI-0. In the second dataset, MCD-C performed better than BCI-0 on all fit metrics. MCD-C+L performed comparably to MCD-C, outperforming BCI-0 on all fit metrics apart from BIC in the first dataset, but it failed to improve the fit relative to MCD-C in a way that would justify inclusion of the dedicated lag parameter.

MCD predictions matched the observed data near-perfectly, which raises the concern that the high flexibility of the MCD model might make it sufficiently adaptable to fit any observed data as long as they roughly follow the shape of a psychophysical curve. This could lead to criticisms that the explanatory heavy lifting is done by simply tweaking time constant values to maximize the model fit. Indeed, response probabilities predicted by MCD models tend to show remarkably high correlations with observed data, as demonstrated not only in the present study but also across multiple simulations of audiovisual spatiotemporal integration (Parise & Ernst, 2016, 2025). To address this overfitting risk, we included a leave-one-asynchrony-out cross-validation procedure in Simulation 1, in which each asynchrony level was predicted by a model fitted to the remaining levels. In both datasets, MCD-C predicted the omitted asynchrony levels better than BCI-0, and its out-of-sample predictions preserved the direction of the asymmetry, including its reversal between the datasets. These results show that high flexibility of the MCD model does not come at the cost of compromised generalizability: the model did not merely reproduce the asymmetry of RHI detection probabilities, but predicted it from incomplete data.

The two model families conceptualize asymmetries observed in RHI probability data in fundamentally different ways. From the perspective of the MCD, the reversed skew reflects individual differences in unisensory processing speeds between the two samples. Participants in the first dataset appear to have processed tactile signals faster, whereas the second sample was, on average, faster at processing visual signals—differences that translated into opposite relations between the tactile and visual time constants estimated for each dataset. The BCI model, which enforces a symmetric distribution of RHI probability, cannot represent asymmetry in either direction and therefore treats the observed skew as measurement noise. The same structural constraint explains why, in cross-validation, BCI-0 mispredicted the omitted asynchrony levels in the direction opposite to the observed asymmetry: the prediction for an omitted level was bound to the predicted RHI probability at the mirror asynchrony with an opposite sign. Nevertheless, cross-validation along a single stimulus dimension remains a limited test, because the general shape of the psychometric function across asynchrony levels is known in advance. A more stringent test would be to predict responses to rich, multidimensional, and unpredictable stimuli, whose variability cannot be reduced to a single scalar parameter such as asynchrony. For example, in Parise and Ernst (2016), the MCD classified independently generated random sequences of flashes and clicks in close agreement with human observers, with predictions following directly from the MCD outputs, without a fitted GLM or any prior specification of the stimulus features driving either human or model classifications.

Simulation 1 also compared two read-outs of the MCD outputs. Adding *MCD_lag_*, which encodes the temporal order of the signals, to the GLM barely improved the fit in either dataset and did not generalize reliably in cross-validation. In our simulations, RHI detection probabilities were captured by *MCD_corr_* alone, with the asymmetries arising from differences between the unisensory time constants. In this sense, MCD-C reproduced RHI detection probabilities that are commonly explained as the outcome of causal inference (Chancel et al., 2022a) on the basis of correlation detection alone, without temporal order information. These results do not rule out a contribution of temporal order processing to body ownership, but indicate that, within the MCD framework, it is not required to account for RHI detection probabilities across asynchrony levels.

The results of Simulation 2 showed that the MCD model predicts stronger visuo-tactile integration—and thus higher RHI occurrence—for irregular and unpredictable tapping patterns, regardless of which of the two datasets the model parameters were fitted to. Two information-theoretic measures were used to evaluate signal complexity: (1) first-order entropy, which was estimated based on ISI variability and thus related to the general irregularity of the signals, and (2) second-order entropy, which was estimated based on variability in ISI pairs and thus related to the absence of periodicity in the signals. Multiple regression analyses revealed positive linear relationships between predicted RHI probability and both signal irregularity and signal unpredictability. The two measures uniquely accounted for comparable shares of variance in predicted RHI probabilities, suggesting that the MCD model predicts both to be relevant for visuo-tactile correlation detection.

In the final Simulation 3, we proposed novel RHI detection metrics that were drawn by analogy from their counterparts in simultaneity detection research: (1) Point of Strongest Ownership, related to the highest expected probability of RHI occurrence, and (2) Window of Subjective Ownership, related to the window of visuo-tactile asynchrony that is sufficiently small for the RHI to reliably occur. The model generated systematic variability in PSO and WSO values, as well as asymmetries in RHI detection probability distributions, by linking predicted RHI probabilities to the time constants of early-stage filters determining visuo-tactile processing times. Lengthening the tactile time constant shifted the PSO toward tactile-first asynchronies and also widened the WSO, whereas jointly lengthening both early time constants widened the WSO with little change in the PSO. Within the MCD framework, the WSO width thus reflects the overall duration of unisensory processing, and the PSO reflects the relative processing speed of the two modalities. This yields the testable prediction that individual differences in RHI susceptibility under different asynchrony conditions may reflect differences in how quickly visual and tactile signals are processed before feeding into the integration stage.

Together, the results of Simulation 2 and Simulation 3 suggest that the MCD model can capture nuanced temporal aspects of visuo-tactile integration for body ownership, accounting for several factors potentially relevant to RHI that may not be readily explained within the Bayesian framework (Litwin, 2020). For instance, the observation that irregular and unpredictable stimulation patterns elicit more vivid RHI was noted in the earliest experimental studies on the RHI (Armel & Ramachandran, 2003; Ehrsson et al., 2005) and later demonstrated empirically (Litwin et al., 2020), but a formal computational account of this effect has not been proposed. This complexity effect cannot be accounted for by the current BCI models, for which performance in the RHI detection task depends only on mean asynchrony values, regardless of the exact temporal structure of the signals. Although a Bayesian observer could in principle account for signal complexity, it would have to be explicitly represented in the model as a dedicated parameter, which requires knowing in advance which stimulus features are relevant. On the other hand, signal complexity effects can emerge naturally from a simple mechanism of correlation detection in the MCD unit. Because the function linking the *MCD_corr_*output to RHI detection probability is monotonic—with higher *MCD_corr_* directly translating to higher predicted RHI probability—the positive linear relation between signal complexity and RHI probability must arise at the level of MCD computations (i.e., how *MCD_corr_*output is calculated by the MCD unit). More irregular sequences contain more clusters of short ISIs, whose overlapping filtered responses amplify each other at the multiplicative stage of the MCD unit, yielding a higher time-averaged *MCD_corr_* output for the same number of stimuli. Therefore, this effect stems from a low-level computational property of the MCD architecture rather than being a model representation of complexity per se.

In the current simulations, the predicted impact of signal complexity was substantial, as it accounted for over 20% of the variance in MCD-predicted RHI probabilities when the relative shift of visuo-tactile signals and stimulus presentation order were fixed. Notably, the MCD model predicted both simple ISI variability and temporal unpredictability to influence visuo-tactile integration, with comparable unique contributions of the two factors within a single regression model. This novel model prediction could be evaluated in future psychophysical studies by independently manipulating first- and second-order entropy of visuo-tactile signals. RHI detection probabilities could be tested under different asynchrony levels, with the influence of signal complexity expected to be strongest at asynchrony values close to the boundaries of individual participants’ WSOs, where uncertainty regarding whether signals should be integrated or segregated is the highest.

Another notable strength of the MCD model is that it conceives of individual differences in RHI susceptibility as idiosyncrasies in visuo-tactile signal processing. Time constants for low-pass filters directly correspond to visuo-tactile processing times, which in turn determine the width and symmetry of RHI detection probability distributions. This offers a candidate mechanistic account of PSO and WSO as consequences of stimulus processing times, which, if successfully tested against individual-level data, could represent an important advantage over Bayesian models. In the BCI model (Chancel et al., 2022a), individual WSOs could potentially be accounted for by introducing an additional free parameter, dedicated to the observer-specific belief about the variability of asynchrony between tactile and visual events. Lower values of this uncertainty parameter should make the model more stringent about what constitutes evidence for a common cause, decreasing integration at larger asynchrony values, and increasing RHI detection at near-zero asynchronies. In this sense, values of this additional uncertainty parameter could thus be proportional to WSO with lower parameter values related to narrower WSOs. However, a previous study (Chancel et al., 2022a) has shown that introducing the parameter to the BCI model does not compensate for the loss in parsimony, as it fails to improve the relative fit. Therefore, although structurally possible, an attempt at accounting for WSO within the BCI framework does not yield clear explanatory benefits.

The MCD model might offer a promising stimulus-driven alternative. Recently, a common neural mechanism for integrating visuo-tactile signals for simultaneity perception and body ownership was suggested (D’Angelo et al., 2026), and MCD might describe the computational underpinnings of this common mechanism based on temporal filtering of sensory signals. In line with MCD predictions, visuo-tactile simultaneity judgments highly correlate with RHI detection judgments (Chancel et al., 2022a; D’Angelo et al., 2026), and visuo-tactile temporal binding window width has been shown to be predictive of the susceptibility to RHI under moderate asynchrony conditions (Costantini et al., 2016). Again, further experimental work is needed to test the specific prediction that simultaneity judgment and RHI detection performance are interconnected as they rely on the same time constant parameters of visual and tactile filters. This could be accomplished by fitting the MCD model parameters to individual participant data in a simultaneity judgment task, and then using these independently estimated temporal constants to predict performance in RHI detection task. Notably, experimental methods allowing determination of asymmetrical visuo-tactile temporal binding windows are already available (Huntley et al., 2023).

The MCD model for temporal integration as presented above does not come without some structural limitations. In our paper, the original MCD model was deliberately chosen as strictly focusing on temporal aspects of integration, mirroring the scope of the BCI model.

This model version consists of a single processing unit that is capable of processing high-frequency signal inputs, but that does not process any spatial information. This is a limitation relative to Bayesian models which allow explicit modeling of spatial properties of multisensory stimulation, including hand location (Fang et al., 2019; Samad et al., 2015) as well as visual and proprioceptive noise (Chancel & Ehrsson, 2023; Chancel et al., 2022a). However, recent developments in MCD architecture have introduced mechanisms for spatial integration that are sensitive to the location and reliability of spatial cues (Parise, 2025).

These newer MCD formulations consist of populations of multiple MCD units organized on a 2D grid, isomorphic to the pixel-wise organization of a visual display, with each unit processing intensity signals from corresponding ‘pixels’. Because a grid of MCD units can process spatial information directly as a two-dimensional signal input—including noise, if present—this architecture could be used to validate the framework in the visual noise regime. Using extended reality setups similar to those in Chancel et al. (2022a), one could compare how MCD and BCI models perform under different noise conditions in a fair modeling context allowing the simulated noise to be presented in forms appropriate for each model (BCI: as a parameter; MCD: as a noisy 2D signal). Another limitation is that the MCD unit itself is insensitive to higher-level factors and does not account for how these factors influence the final multisensory integration outcome. Even though the full MCD modeling pipeline effectively incorporates a decisional layer in the form of converting *MCD_corr_*output into response probabilities via the GLM*—*which captures certain cognitive biases in a way similar to Bayesian priors (Parise, 2025)*—*such statistical inferences might not be sufficient to explain the full scope of higher-level influences on body ownership. For instance, the discrepancy between the widths of temporal binding windows and WSOs could be explained by introducing a coupling prior that promotes the sense of ownership through the assumption that spatially congruent visual and tactile signals tend to originate from the same location (Litwin, 2020). These priors might include information about the expected size of the area of possible tactile interactions with the world—the peripersonal space (Noel et al., 2018). Empirical evidence suggests that spatial integration for RHI may be governed by spatial coupling priors rather than linear distance: RHI strength sharply declines for distances larger than 30 cm (Kalckert & Ehrsson, 2014; Kalckert et al., 2019), but not necessarily for smaller distances (Motyka & Litwin, 2019), especially when classic RHI paradigm and complex tactile stimulation are used (Litwin et al., 2020). This suggests that when the distance between hands exceeds 30 cm, it is *a priori* too large to promote visuo-tactile fusion, because the rubber hand lies outside the peripersonal space. Such causal inferences incorporating nuanced prior information may pose an explanatory challenge for the MCD framework.

The final concern—although not strictly a limitation yet—pertains to the generalizability of MCD to slowly unfolding dynamics of body ownership illusions. MCD was originally designed to detect correlations over very short timescales (1 second; Parise & Ernst, 2016), while illusion elicitation in RHI detection paradigm requires sustained tapping for at least 10 seconds (Chancel & Ehrsson, 2020), and RHI onset is only this rapid after preselection for RHI prone participants (Riemer et al., 2019). Accordingly, signals generated in our study lasted 36 seconds, representing a further extension into an entirely different temporal regime compared to all previous MCD applications. The neural mechanisms required to sustain correlation detection at such longer timescales might differ from short-term correlation detection, as they could include additional processes such as evidence accumulation (Bizley et al., 2016). This is particularly relevant in the context of the temporal dynamics of RHI, since RHI strength rises continuously for at least 2 minutes with synchronous stimulation (Finotti et al., 2023). Lanfranco et al. (2025) provided direct empirical evidence for this gradual build-up, using a drift–diffusion model to show that the RHI emerges and gradually strengthens through the accumulation of multisensory evidence over repeated visuotactile stimulation. It is unclear whether such accumulation effects could arise from sustained correlation detection dynamics; this question should be addressed by an in-depth analysis of the computational structure of the MCD.

Further work is also needed to answer outstanding questions regarding MCD implementation in the brain. For example, although the MCD framework might be better characterized as an account of front-end multisensory processing architecture (Parise, 2025), it is not entirely clear which neural processing stage corresponds to the MCD output. For the modeling purposes, we assumed that *MCD_corr_* represents an all-encompassing correlation estimate, but further central operations on the MCD unit outputs—such as recalibration mechanisms compensating for differences in temporal processing of visual and tactile signals, correcting asymmetries—are certainly possible. From this perspective, MCD and BCI approaches could perhaps complement each other and be integrated, with MCD serving as a lower-level correlation detection mechanism feeding into BCI at higher processing levels.

However, determining common ground and resolving all theoretical differences between the two frameworks was beyond the scope of the present study, which aimed to show that a stimulus-computable correlation detector can comprehensively account for temporal visuo-tactile integration underlying RHI detection.

## Conclusions

In the current study, we applied the MCD framework for multisensory integration (Parise & Ernst, 2016) to model RHI occurrence probabilities under different asynchrony conditions as a function of temporal visuo-tactile correlations. This modeling strategy offers a promising alternative to existing computational approaches, with several unique strengths. The MCD model for RHI not only accounts for observed probabilities in the RHI detection task, but also offers a mechanism by which stochastic visuo-tactile stimulation leads to an increased probability of RHI occurrence, which is in line with empirical findings (Litwin et al., 2020). The model also captures asymmetries in response probabilities across varying visuo-tactile asynchronies, generating the prediction that susceptibility to RHI under temporal mismatch may be linked to individual differences in visuo-tactile processing speeds, and thus to temporal binding window widths (Costantini et al., 2016). Importantly, the model is biologically inspired and computationally plausible (Parise & Ernst, 2016). While further experimental and neuroscientific validation is needed, the present study provides support for the possibility that the computational mechanism underlying body ownership may depend on detecting multisensory correlations in a manner analogous to motion perception.

## Additional information

## Author contributions

Piotr Litwin is the sole author of the manuscript.

## Data availability

No data was gathered for the purpose of this study. The original databases used in this study are available at https://osf.io/n7atw/ and https://osf.io/qjf5h.

The MATLAB code for all simulations, analyses, and figures is available at https://osf.io/cy7k5/ under the MIT License.

## Author Note

This research was funded by the National Science Centre, Poland (NCN) (Grant 2024/55/D/HS6/00965). I have no conflict of interest to disclose. No data was gathered for the purpose of this study. The original databases used in this study are available at https://osf.io/n7atw/ and https://osf.io/qjf5h. The code for the data simulation and analysis as well as supplementary material are available at https://osf.io/cy7k5/. Correspondence concerning this article should be addressed to the Faculty of Psychology, University of Warsaw, Banacha 2D, 02-097, Warsaw, Poland.

