## Appendix 1 and Supplementary Figures S1-S4 for "The Multisensory Correlation Detector model explains temporal integration in the Rubber Hand Illusion"

#### Appendix 1 - key MCD equations

The Multisensory Correlation Detector (MCD) is a biologically plausible model for multisensory integration, inspired by the Hassenstein–Reichardt elementary motion detector. In the MCD model for RHI, the MCD unit receives time-varying visual and tactile input signals and produces two outputs: a correlation signal ($MCD_{corr}$) encoding the strength of the temporal cross-correlation, and a lag signal ($MCD_{lag}$) encoding the temporal order between the inputs.

Signal processing within the MCD unit consists of two stages. In the modality-specific filtering stage, visual and tactile signals are independently convolved with a modality-specific low-pass temporal filter. These filters are modeled as exponential functions:

$$(1)\ f_{mod}(t) \ = \ t \cdot exp(- \frac{t}{\tau_{mod}})$$

where $\tau_{mod}$ is the modality-dependent time constant ($\tau_V$ for vision, $\tau_T$ for touch), and $t \geq 0$. The unisensory signals are filtered using:

$$(2)\ MCD_V(t) \ = \ f_V(t) * s_V(t)$$

$$(3)\ MCD_T(t) \ = \ f_T(t) * s_T(t)$$

where $s_V(t)$ are visual signals, $s_T(t)$ are tactile signals, and * denotes the convolution operator.

Then, filtered signals enter a multisensory integration stage in which they are independently combined by two mirror-symmetric correlation detectors (subunits). In the first subunit, the additional visuo-tactile low-pass filter (with time constant $\tau_{VT}$) is applied to the visual signal before multiplying it with the tactile signal. In the second subunit, the same cross-modal low-pass filter with an identical time constant $\tau_{VT}$ is applied to the tactile signal before multiplying it with the visual signal:

$$(4)\ MCD_{S1}(t) \ = \ [MCD_V(t) * f_{VT}(t)] \cdot MCD_T(t)$$

$$(5)\ MCD_{S2}(t) \ = \ [MCD_T(t) * f_{VT}(t)] \cdot MCD_V(t)$$

The correlation output is obtained by multiplying the responses of the two subunits:

$$(6)\ MCD_{corr}(t) \ = \ MCD_{S1}(t) \cdot MCD_{S2}(t)$$

$MCD_{corr}(t)$ is a time-varying signal whose magnitude reflects the strength of the temporal cross-correlation between the visual and tactile inputs.

The lag output is obtained by combining the subunit responses through addition:

$$(7)\ MCD_{lag}(t) \ = \ -MCD_{S1}(t) + MCD_{S2}(t)$$

$MCD_{lag}(t)$ is a time-varying signal whose sign indicates the temporal order of the inputs (i.e., which modality leads), and whose magnitude reflects the size of the lag.

To reduce the time-varying outputs to scalar decision variables (one value per trial), the outputs are averaged over an integration window of duration T. These time-averaged values serve as predictors for behavioral responses: $MCD_{corr}$ predicts simultaneity/causality/common cause judgments, while $MCD_{lag}$ predicts temporal order judgments.

The time-averaged outputs are related to response probabilities through a generalized linear model with a probit link function. In the present study, $MCD_{corr}$ (averaged over input signal duration) was used as a predictor of RHI detection (i.e., common cause judgment) probability, either alone (MCD-C) or together with the time-averaged lag output as a second predictor (MCD-C+L):

$$(8)\ P(RHI) \ = \ \Phi\,(\beta_0 + \beta_1 \cdot \overline{MCD_{corr}})$$

$$(9)\ P(RHI) \ = \ \Phi\,(\beta_0 + \beta_1 \cdot \overline{MCD_{corr}} + \beta_2 \cdot \overline{MCD_{lag}})$$

### Supplementary Material

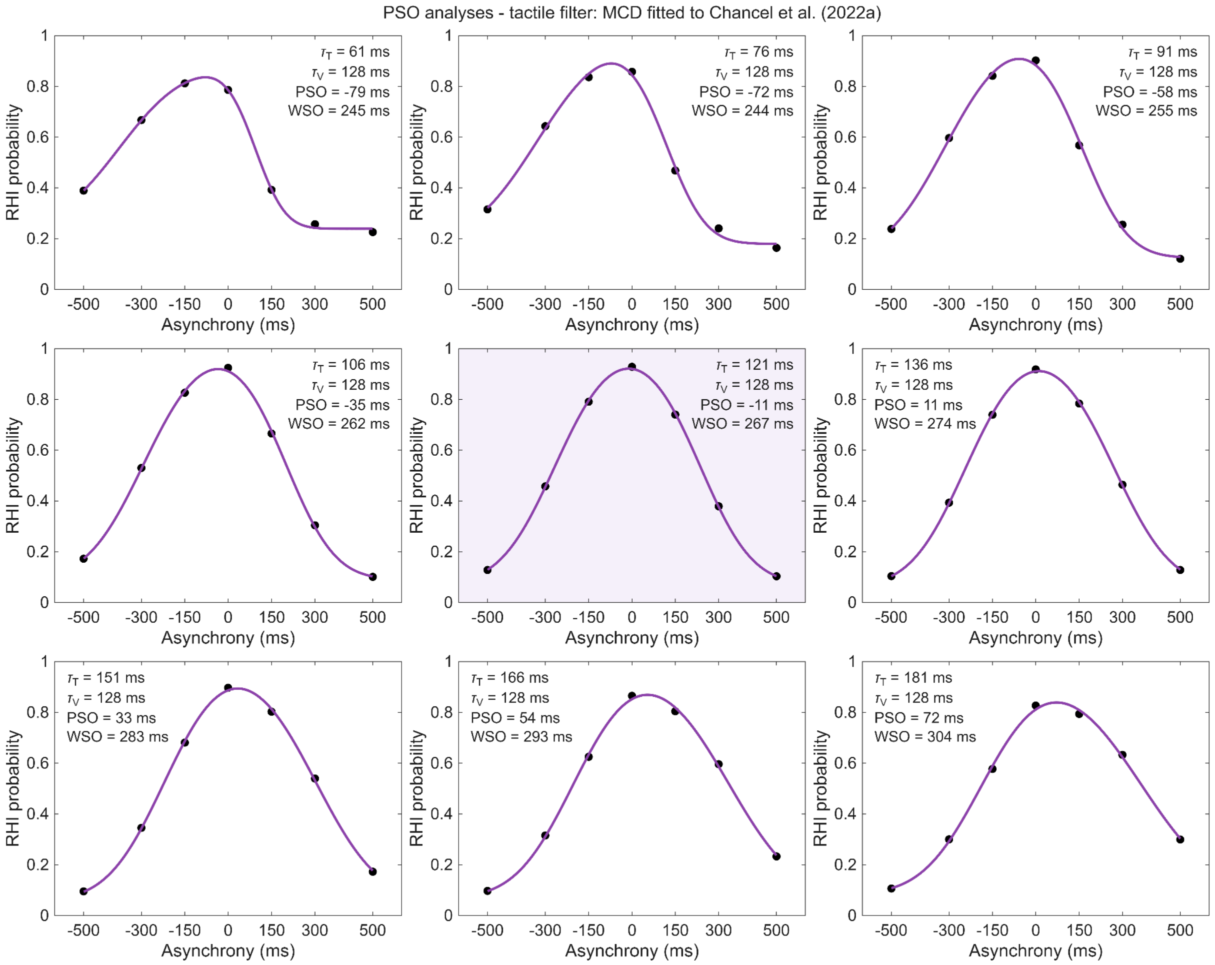


Figure S1. Predicted RHI probabilities for all nine values of the tactile time constant: MCD-C fitted to Chancel et al. (2022a). The tactile time constant was varied in 15 ms steps from 60 ms below to 60 ms above its baseline value (shaded panel: parameter set from Simulation 1); the visual and late-stage time constants were kept at their baseline values. For each parameter set, the GLM read-out was fitted again to the observed RHI probabilities of the dataset. Black dots represent MCD-predicted RHI probabilities and purple lines the fitted asymmetric psychometric functions. Values in each panel give the early filter time constants and the resulting PSO and WSO.

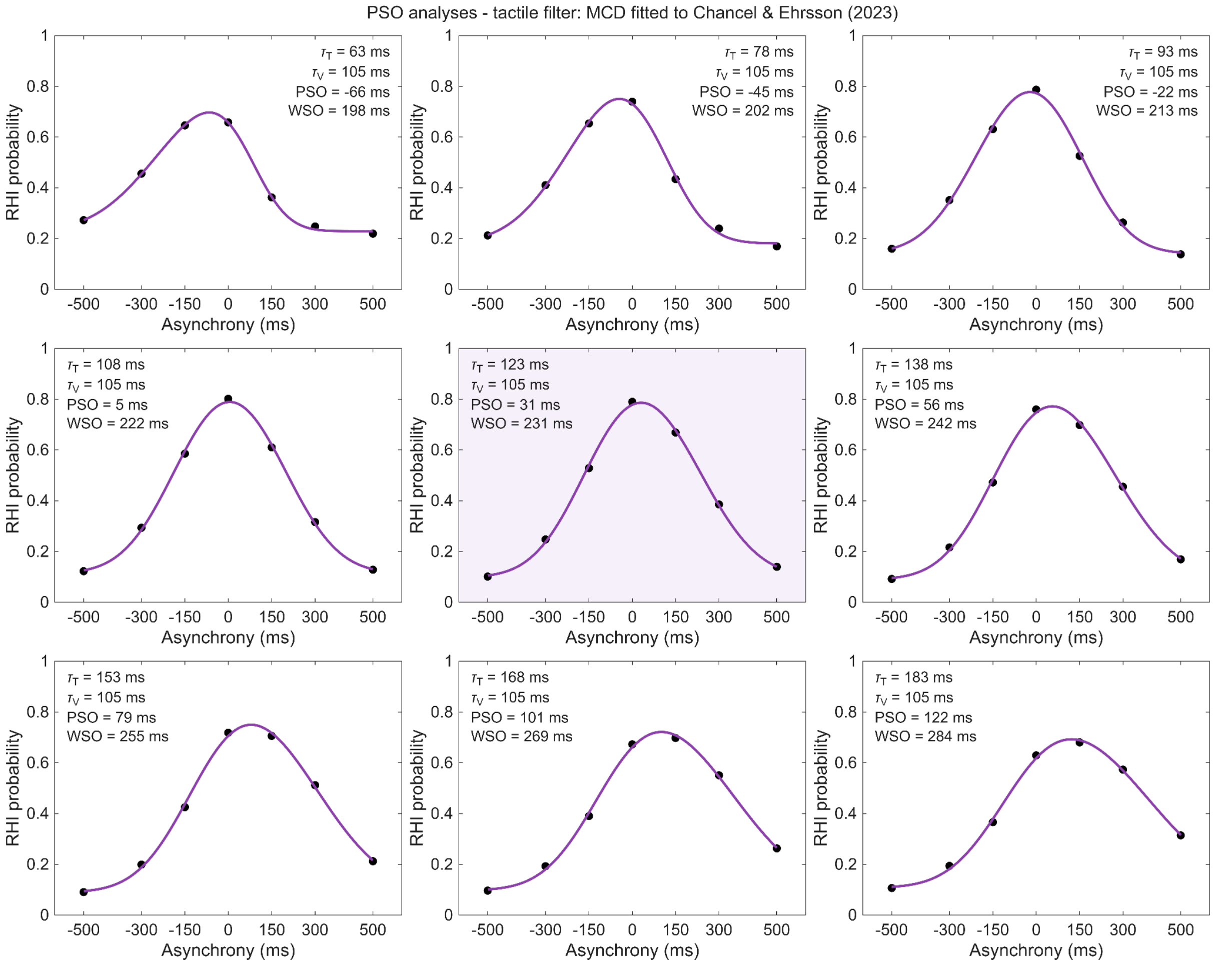


Figure S2. Predicted RHI probabilities for all nine values of the tactile time constant: MCD-C fitted to Chancel and Ehrsson (2023). Manipulation, layout, and symbols as in Figure S1.

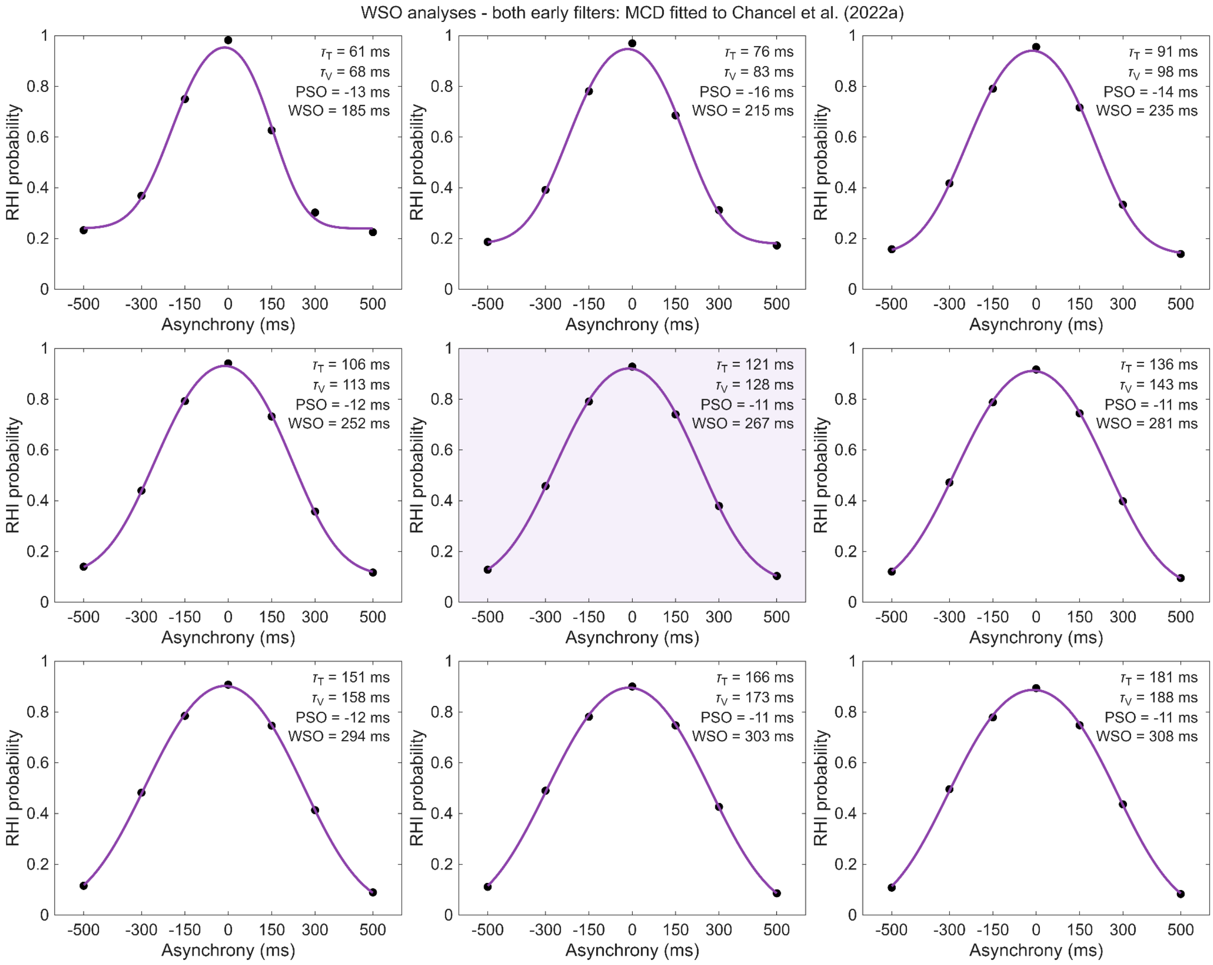


Figure S3. Predicted RHI probabilities for all nine joint values of the early time constants: MCD-C fitted to Chancel et al. (2022a). The tactile and visual time constants were jointly varied in 15 ms steps from 60 ms below to 60 ms above their baseline values (shaded panel: parameter set from Simulation 1); the late-stage time constant was kept at its baseline value. For each parameter set, the GLM read-out was fitted again to the observed RHI probabilities of the dataset. Black dots represent MCD-predicted RHI probabilities and purple lines the fitted asymmetric psychometric functions. Values in each panel give the early filter time constants and the resulting PSO and WSO.

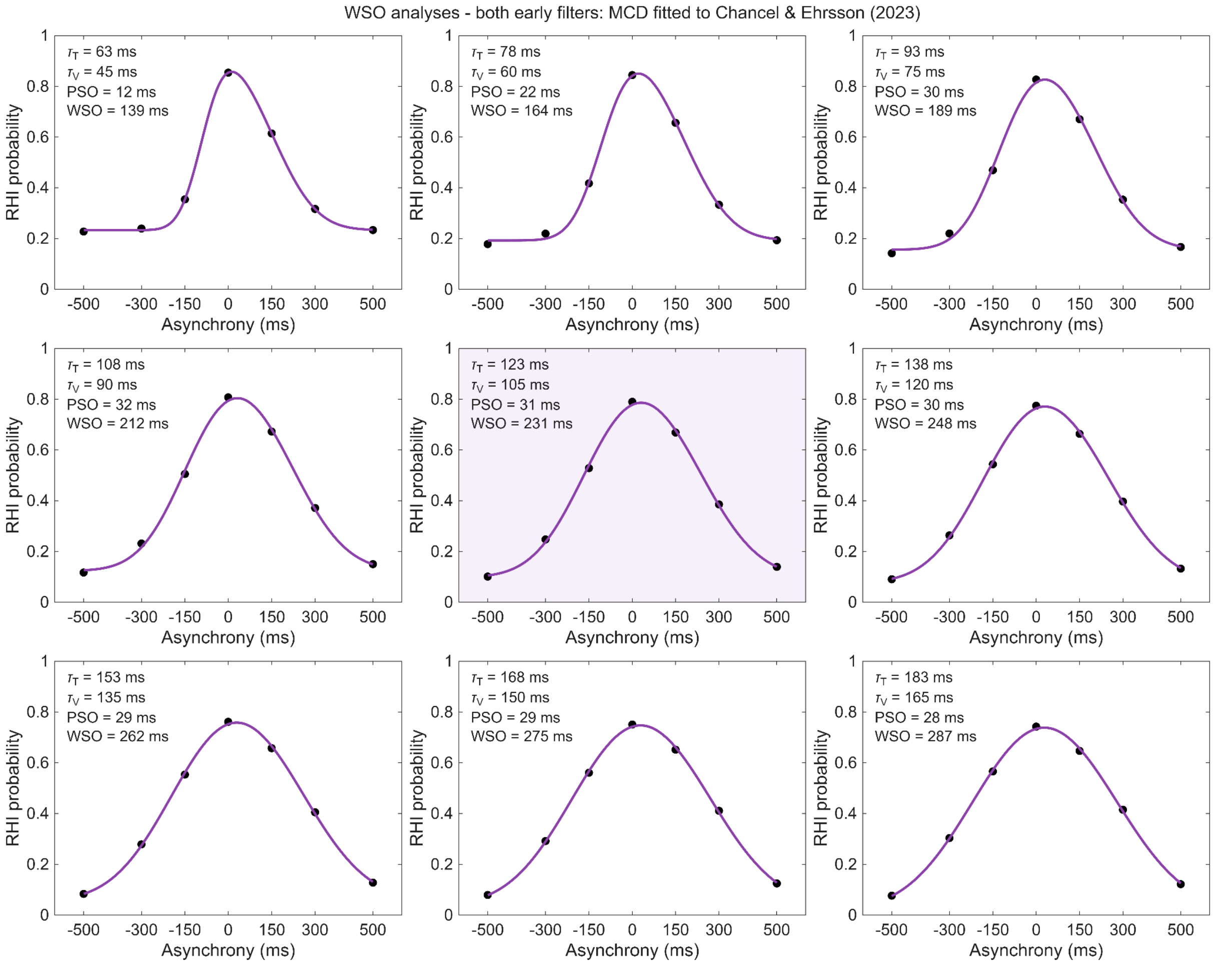


Figure S4. Predicted RHI probabilities for all nine joint values of the early time constants: MCD-C fitted to Chancel and Ehrsson (2023). Manipulation, layout, and symbols as in Figure S3.
